# Functional characterization of dynamic nascent RNA folding ensembles in real-time

**DOI:** 10.1101/2025.10.23.684182

**Authors:** Kavan Gor, Eva Maria Geissen, Olivier Duss

## Abstract

RNA structure starts forming co-transcriptionally as the nascent RNA emerges from the RNA polymerase and is dynamically modulated by cellular factors. How individual RNA conformations, out of an ensemble of RNA molecules, relate to function is not well understood. Here, developing multi-color single-molecule fluorescence microscopy experiments, we track in real-time nascent RNA structure formation, functionally characterizing up to 8 different types of RNA molecules. We find that ribosomal proteins, RNA modification enzymes or antisense oligonucleotides specifically modulate a subset of the RNA folding classes. For example, we provide direct evidence that increased local RNA accessibility at specific sites correlates with the chaperoning activity of ribosomal proteins during ribosome assembly. These experiments provide a general framework to study how dynamic RNA folding, and misfolding, relates to function.

## Main text

RNA structure is central to multiple cellular processes across the tree of life. The ability of RNA to form complex structures enables the binding of various cellular factors such as proteins, nucleic acids, ions and small molecules which can together undergo dynamic conformational changes to impart function (*1-4*). Despite the indispensable role of structured RNAs, the exact rules governing the process of RNA folding that leads to formation of complex functional structures remain elusive, mainly due to RNAs dynamic and flexible nature, often assuming multiple conformations (*5, 6*). These properties also explain the lower performance of deep learning methods for RNA structure prediction in comparison to proteins (*7-10*).

Advances in computational methods to analyze high-throughput RNA structural mapping data have allowed the dissection of RNA structural heterogeneity by providing single-molecule snapshots of RNA structure in cellulo (*11-14*). However, these methods do not provide real-time dynamic information and thus, make direct correlation of RNA structure to function difficult. Furthermore, the complex cellular environment complicates examining the specific effects of various factors on RNA structure formation, making it difficult to learn the biophysical principles of how RNA folds, because the majority of RNAs are expected to “fold correctly” in vivo.

In contrast, in vitro studies allow dissecting the effect of various factors affecting RNA folding, one at a time. For instance, folding intermediates of various RNAs have been previously studied by DNA oligo hybridization to specific sites of RNA, followed by RNase H cleavage, reporting on single-stranded RNA regions (*15*). A similar approach was employed to study the role of RNAP pausing in the co-transcriptional folding of the RNase P RNA, the SRP RNA and the tmRNA, thereby highlighting that the evolutionarily conserved pause sites enable formation of non-native labile RNA secondary structures that sequester the upstream regions of the RNA until the complementary strands are transcribed to form long-range helices (*16*). While these studies have provided an initial framework to assess the kinetics of RNA folding, they did not dissect the RNA folding heterogeneity nor provided real-time dynamic information.

In contrast, state-of-the-art single molecule studies have probed in real-time dynamic RNA folding of small pre-transcribed RNAs, such as riboswitches (*17-21*) or segments of ribosomal RNA (rRNA) (*2, 22*), investigated the effect of transcription on RNA folding (*23-29*), and used protein binding as an indirect readout for RNA folding efficiency (*24, 28*) but have not directly correlated dynamic nascent RNA structure formation with function at the single RNA level. For example, Chauvier et. al have used the concept of dynamic RNA structure probing (SiM-KARTS) using short DNA probe binding to the RNA to study the folding kinetics of the fluorine riboswitch (*23*). Their assays highlight the dynamic competition between the fluorine ligand and the NusA protein, which governs transcription regulation, but their approach does not track RNA structure and protein binding simultaneously. Furthermore, such studies have been performed only on relatively small RNA molecules, which effectively reduces the probability of RNA molecules to misfold.

Large, highly structured RNAs like the 16S rRNA fold into complex secondary and tertiary structures co-transcriptionally, bind multiple r-proteins sequentially, and are chemically modified and processed simultaneously to eventually form ribosomes (*30-33*). Early studies have investigated the vectorial (5’ to 3’) folding of rRNA and the sequential as well as cooperative nature of r-protein assembly (*34-39*). Technological advancements allowed probing reaction kinetics that highlighted the presence of parallel and heterogeneous folding pathways (*40-44*). While these studies used pre-transcribed and pre-folded RNA and correlated the results into a co-transcriptional folding model, recent studies using single-molecule fluorescence microscopy on nascent RNA have directly visualized the co-transcriptional aspect of r-protein assembly (*24, 25, 28, 45*). These studies have pointed out the heterogeneity of the assembly process and associated it with RNA misfolding, but have only partially investigated the folding dynamics of the RNA directly (see supplementary text) (*25*).

Here, using multicolor single-molecule fluorescence microscopy, we are simultaneously tracking transcription elongation of the 3’ domain of the 16S rRNA, are probing the accessibilities of RNA sites on the nascent RNA, which includes co-transcriptional as well as post-transcriptional RNA structure relaxation, and are assessing the binding of a protein as a readout for functional RNA structure formation. Using up to 5 different fluorescent dyes within a single experiment, we can simultaneously study the dynamics of up to 8 different nascent RNA molecule types (folding classes). We see how various factors, such as RNA modification enzymes, ribosomal proteins (r-proteins) or antisense oligonucleotides (ASOs), specifically affect a subset of these folding classes. Our assays illustrate a variety of misfolding behaviors and dissect the heterogeneity of the structural ensemble during ribosome assembly. Overall, we provide an unprecedented view of the complex and understudied process of dynamic RNA folding and misfolding.

## Results

### Short DNA probes can be used to detect dynamic RNA accessibility profiles

In order to provide dynamic structural information on single RNA molecules in real-time, we employed a previously established concept in which the transient binding of short DNA probes to specific RNA sites serves as a proxy for single-stranded RNA at these sites (*19, 46*) (Fig. 1). As positive controls for RNAs with maximal probe accessibility, we first tracked the binding of short 7 nucleotides (nt) long DNA probes to short single-stranded target RNAs (ssRNAs) that were immobilized to a glass surface for single-molecule imaging by total internal reflection fluorescence (TIRF) microscopy (Fig. 1A). Specific and transient binding of the short DNA probe to the ssRNA was detected by FRET between the ssRNA, labeled with a donor dye (Cy3), and the DNA probe, labeled with an acceptor dye (Cy5, Cy5.5 or Cy7). We chose three different single-stranded RNAs comprising short stretches (7 nt) of the 3’domain of the *E. coli* 16S rRNA: 1) the linker region between helix 28 (H28) and H29 (residues 933-939; in the following referred as to H2829), which is part of the extended binding site for primary binding r-protein S7, 2) the 5’half of the tip of long-range H30 (950-956), which needs to form to stabilize the S7 binding site, and 3) the 5’-half of H32 (986-992), a long-range helix, which is not part of the S7 binding site (Fig. 2A, fig. S1A) (*34*).

**Fig. 1:**
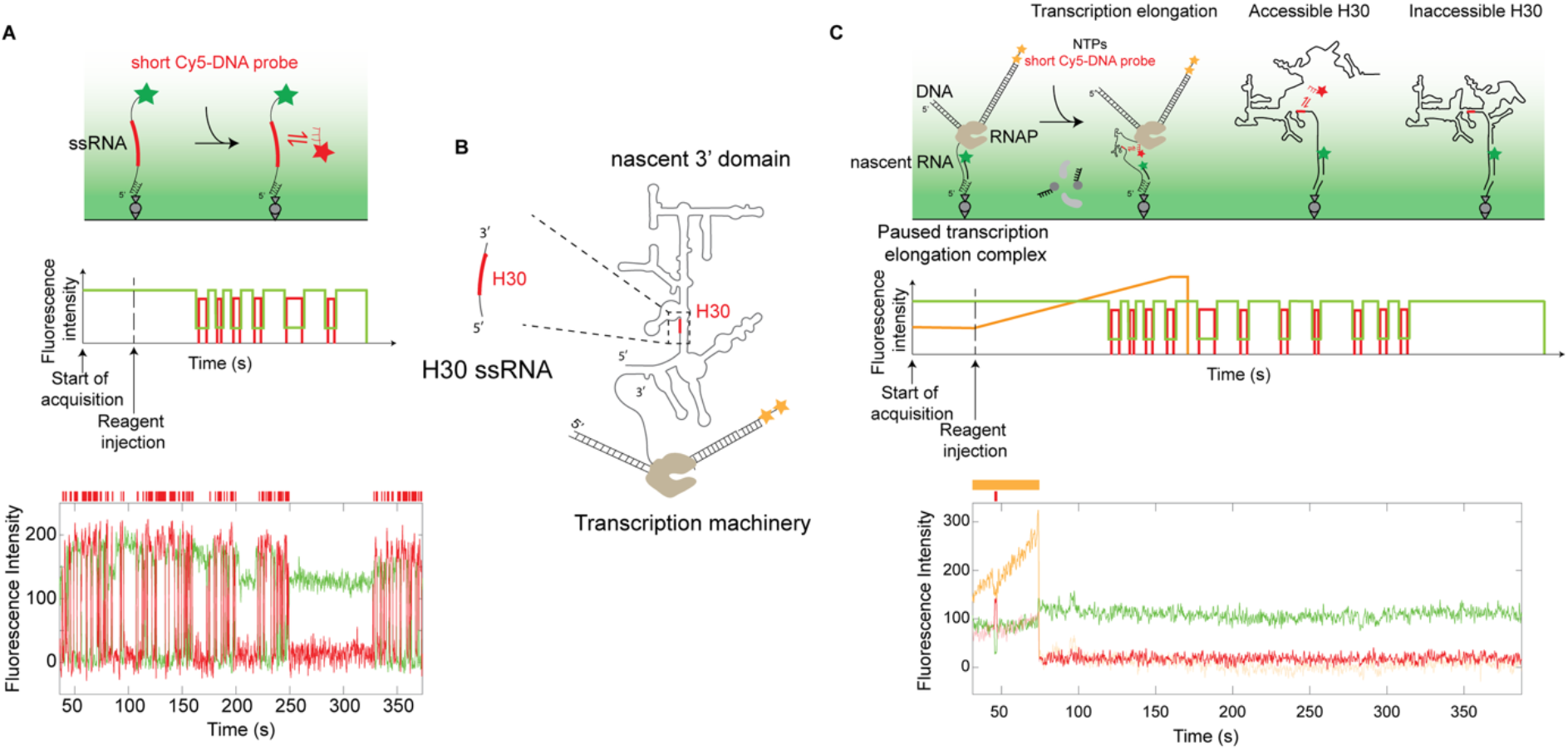
Concept of single-molecule dynamic RNA structure probing assay. **(A)** Schematic of experimental setup to probe single-stranded RNA (ssRNA) (top), idealized trace (center), and an example smoothed trace (bottom) showing the binding and unbinding of the probe (red – acceptor Cy5 dye) to the ssRNA (green – donor Cy3 dye) detected using FRET. **(B)** Schematic of H30 ssRNA (left; used in **(A)**) and nascent 3’ domain transcribed by the transcription machinery (right; used in **(C)**). **(C)** Schematic of the experimental setup to simultaneously monitor transcription and RNA accessibility in real-time (top), idealized trace (center), an example smoothed trace (bottom) showing transcription (yellow – Cy3.5 dye) and H30 probe binding (red – acceptor Cy5 dye and green – donor Cy3 dye).

**Fig. 2:**
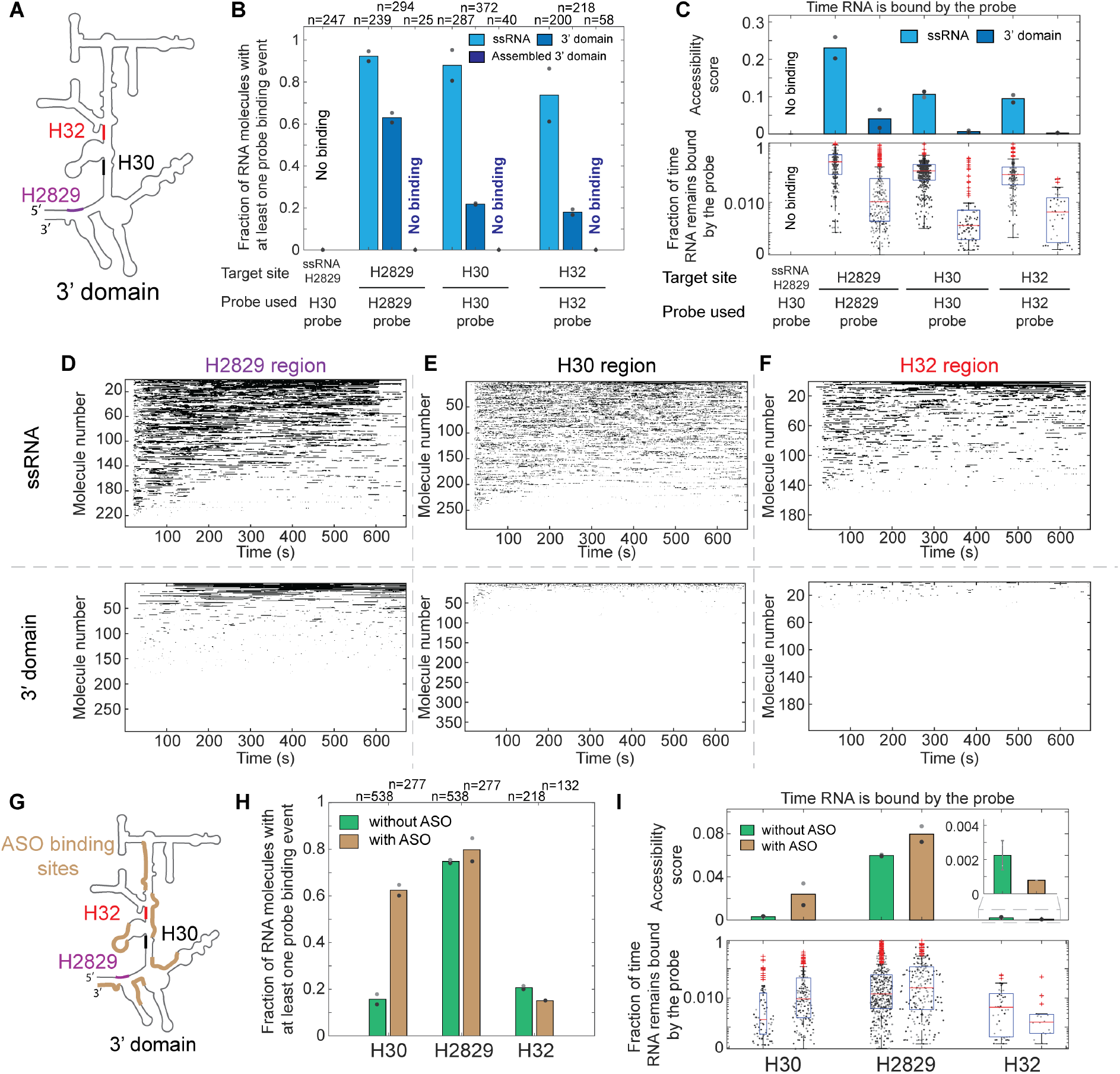
Monitoring RNA accessibility of helices in the 3’domain of the 16S rRNA. **(A)** Schematic of the 3’domain highlighting the different regions probed. **(B)** Fraction of RNA molecules accessible by DNA probes. **(C)** The accessibility score is the mean fraction of time the RNA molecules remain bound by the probe (top). The fraction of total experimental time the individual RNA molecules remain bound by the probe is represented as individual dots (bottom plots). **(D-F)** Rasterplot representation showing individual molecules as rows with black and white bars representing probe bound or unbound, respectively. **(G)** Schematic of the 3’domain with the six ASO binding sites highlighted. **(H**,**I)** Effect of ASOs on the fraction of RNA molecules accessible by the DNA probes **(H)** and RNA accessibility score and the fraction time bound by probe for individual RNA molecules **(I)**. The without ASO data (in **(H**,**I)**) is the same as presented in Fig. 2B. **(B**,**H)** Number of molecules (n) used for analysis is indicated. Same number of molecules (n) were analyzed for **(B)** and **(C)**; and **(H)** and **(I). (C**,**I)** The median and the quartiles in the overlayed boxplot are calculated by pooling the replicates. **(B**,**C**,**H**,**I)** Mean of two replicates is shown as bar, individual replicates shown as light and dark grey dots.

We find that for the specific targets, around 90% of the RNA molecules show repetitive DNA probe binding events, in agreement with a mostly single-stranded RNA structure (Fig. 2B). In contrast, we do not detect probe binding events to a non-target RNA (H30 probe does not detectably bind to H2829 ssRNA) (Fig. 2B and methods). The majority of the DNA probe accessible RNA molecules remain accessible during the entire 10 minutes experimental time, which we quantify by an RNA molecule-specific accessibility score (see methods and Fig. 2C). The on-rate of the probe is linearly dependent on the concentration of the probe as expected for a single fully accessible RNA conformation (concentration independent on-rate: 1.8 × 10^-6^M^-1^s^-1^ for H2829; fig. S1B) (*46-48*). At 200 nM probe concentrations, the median DNA probe arrival times are 3.2 s, 11.5 s, 10.2 s and the median bound-lifetimes of individual binding events of our probes are 1.6 s, 5.7 s and 11.5 s for H30, H32, and H2829, respectively. Kinetic analysis of the bound lifetimes indicates single exponential behavior (fig. S1C). Overall, this setup allows us to track specific RNA accessibilities in a timescale of a few seconds resolution.

### Site-specific RNA accessibilities during folding of the nascent 16S rRNA 3’domain

Next, we sought to investigate how these various RNA segments engage in RNA structure formation in context of an actively transcribing RNA which is highly structured once entirely transcribed and natively folded. We chose the 3’domain of the 16S rRNA in *E. coli* (Fig. 1B) as it was previously well characterized by various biophysical and biochemical approaches. For example, we previously tracked the co-transcriptional formation of the long-range H28 by donor and acceptor dyes located at the 5’-end and 3’-end of the 3’domain (*25*). While these data allowed us to determine whether a single RNA molecule had H28 formed or not, technical limitations (see supplementary text) prevented us from determining at what timepoint this helix was formed and from providing dynamic site-specific RNA folding information on the rest of the > 440 nucleotides long 3’domain intervening RNA.

To this end, we immobilized a stalled transcription elongation complex via the 5’ end of the nascent RNA (see methods) to the surface for multi-color single-molecule imaging (Fig. 1C). At the start of the reaction, we delivered a short DNA probe and NTPs to the immobilized stalled complex, which triggered transcription elongation of the 3’domain rRNA (Fig. 1C). As the transcription progresses, the Cy3.5-labeled 3’ end of the DNA template approaches the imaging surface causing an exponential increase in fluorescent signal intensity, which reports on transcription elongation (*24, 25*). Our labeling strategy allowed us to simultaneously track active transcription elongation, determine at what timepoint the full-length 3’domain RNA has completed transcription (complete signal loss of both Cy3.5 dyes upon dissociation of the DNA template from the surface as opposed to stepwise photobleaching of both Cy3.5 dyes) and when the specific RNA sites are accessible during transcription and following dissociation of the transcription machinery of the nascent RNA (fig. S1D), thus providing site-specific and dynamic RNA structure information of single nascent RNA molecules.

Kinetic characterization of probe binding to the 3’domain shows single-exponential behavior for the regions probed, except for the H2829 region, which has an additional minor population (fig. S1C and supplementary text). RNA mutants disrupting the probe binding target sites show that the probes bind specifically, as we do not detect off-target binding to any part of the entire 3’domain (fig. S2A-E). To ensure that our FRET-based probe detection approach captures all the probe binding events in the different possible RNA conformational states, we repeated our experiments with a Cy3B-Cy5 FRET pair to extend the accessible FRET range (Cy3B-Cy5: R_0_=71.9 Å; Cy3-Cy5: R_0_=51 Å; https://www.fpbase.org/fret/) and find comparable fractions of molecules with probe binding for all three measured sites (fig. S2F and supplementary text). We also verified that the transient binding of our probes does not detectably affect co-transcriptional RNA folding: The fraction of RNA molecules that bind r-protein S7 (proxy for successfully folded RNA molecules; see below) is not significantly affected, as well as the time from transcription initiation till appearance of the first S7 binding event is unaffected by the presence of the DNA probes targeting various RNA sites (fig. S1E,F). Overall, all these controls demonstrate that we can robustly track nascent 3’domain RNA structure formation using our short DNA probes.

In contrast to the short ssRNAs, for the entire 3’domain rRNA, the fractions of nascent RNA molecules that bind the probe substantially decrease for all the sites. They decrease from 80-90% to ∼20% for H30 and H32. In contrast, still ∼60% of the RNA molecules remain accessible to the H2829 junction DNA probe suggesting that local accessibility dynamics vary between different sites on the naked rRNA (Fig. 2A,B,D-F). For the fully assembled 3’domain, identified by bound tertiary r-protein S3-Cy5.5, we did not observe any probe binding (Fig. 2B and fig. S3), in line with the expectation that these sites are completely protected in the natively assembled 3’domain (*34, 37*). When comparing short ssRNAs versus the nascent 3’domain rRNA, the median RNA molecule-specific accessibility score reduced for all the sites by 1-2 orders of magnitude (Fig. 2C), in agreement with global co-transcriptional RNA structure formation (*34, 37, 40, 42, 49*). We additionally observe high variation in this accessibility score between molecules, which we attribute to the heterogeneous conformational nature of RNA molecules (see supplementary text).

Next, we designed ASOs that can bind to different regions of the 3’domain with the aim to perturb co-transcriptional RNA folding dynamics (Fig. 2G). In contrast to the short DNA probes (7 nt) that were designed to bind transiently to the nascent RNA, the DNA ASOs were designed to remain stably bound to the nascent RNA and therefore, had a length of 18 nucleotides (*50*). Upon simultaneous addition of six ASOs covering several regions across the entire 3’domain rRNA, accessibility of the H30 site increases from ∼18% to ∼62%. In contrast, for the H2829 and H32 sites, we do not detect substantial accessibility increases despite the presence of the 6 ASOs (Fig. 2H,I; fig. S4).

Overall, probing the accessibility of different sites across the nascently transcribed 16S rRNA 3’domain, our data show that sites, which are predicted to be inaccessible in the natively folded 16S rRNA structure, are transiently accessible in a subpopulation of RNA molecules, at various degrees for the different sites. This raises the question whether the subset of probe-accessible RNA molecules may constitute misfolded RNA molecules.

### Correlating rRNA accessibility with r-protein binding as functional readout

In order to functionally characterize the heterogenous pool of nascent rRNA conformations at the single RNA level, we repeated our site-specific RNA accessibility probing experiments in presence of r-protein S7 (Fig. 3A). The binding site of primary r-protein S7 extends across and covers the 3-way junction of H28/H29/H43 as well as the 4-way junction of H29/H30/H41/H42 (Fig. 3B and fig. S5A). Thus, successful binding of S7 to the nascent RNA serves as a convenient proxy for assessing the native folding state of the entire lower part of the 3’domain (*25*).

**Fig. 3:**
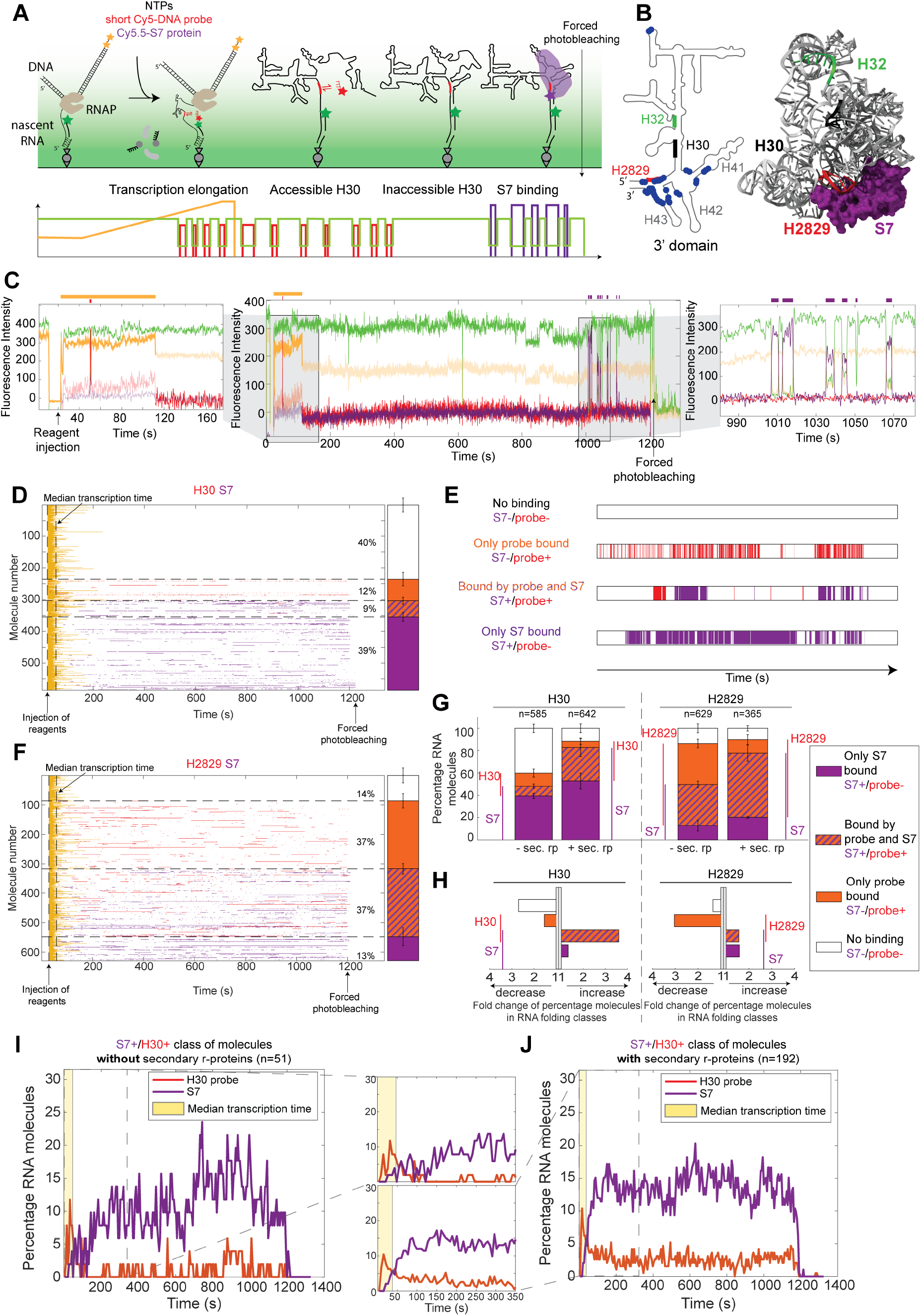
Assigning function to individual RNA folding classes. **(A)** Schematic of the experimental setup. **(B)** Schematic (left) and 3-dimensional structure (right) of the 3’domain highlighting the different regions probed: H2829 (red), H30 (black), H32 (green) and contact sites of S7 to RNA (blue circle) (PDB accession code: 4V9P). **(C)** Idealized trace (top) and an example smoothed trace (bottom) showing transcription (yellow – Cy3.5 dye), FRET donor (green – Cy3 dye), short DNA probe that is probing RNA accessibility (red – Cy5 dye) and binding of r-protein S7 (purple – Cy5.5 dye). **(D**,**F)** Rasterplots depicting the individual molecules as rows with transcription (yellow), DNA probe (red) and S7 (purple) binding events shown as colored bars for H30 **(D)** and H2829 **(F)** sites, respectively. The sorted rasterplots show 4 RNA folding classes: No binding events as S7-/probe-class (white); only probe bound class as S7-/probe+ (orange); S7 bound and probe bound class as S7+/probe+ (stripes of orange and purple); only S7 bound class as S7+/probe-(purple). **(E)** A simplified example trace from each class is shown. **(G**,**H)** Percentage of molecules in RNA folding classes **(G)** and relative fold change **(H)** upon addition of secondary r-proteins (sec. rp). **(I**,**J)** Time dependent analysis of the S7+/H30+ class of molecules in absence **(I)** and presence **(J)** of the secondary r-proteins. **(I**,**J)** The region highlighted in yellow represents the median time RNA is associated to the transcription machinery. **(D**,**F-J)** Data represented by pooling three replicates, error bars show weighted standard deviations, number of molecules analyzed (n) is shown.

In order to simultaneously track site-specific RNA accessibility (DNA probe binding) and functional RNA structure formation (S7 binding to the nascent RNA), we added 20 nM Cy5.5-labeled S7 in addition to Cy5-DNA probe and NTPs to our nascent RNA folding assay (Fig. 3A,C). FRET-based readout allows us to detect only specific S7 binding events, which we additionally verified by mutating the rRNA at its S7 binding site: This abolished the S7 binding almost completely (fig. S5B-D), in line with previous reports that used an S7 mutant to show the same (*25*). The median arrival time of S7 to wildtype 3’domain RNA was ∼12 s, and the bound lifetime could be classified into two average S7-bound phases: short-lived (∼1 s) and longer-lived (∼10-20 s), as described previously and summarized in Table S1 (fig. S5E).

In order to investigate the dynamic RNA conformational heterogeneity, we classified all the RNA molecules into four classes: Two S7 binding-competent classes S7+/probe-(no probe binding) and S7+/probe+ (with probe binding), which we assign as correctly folded and thus functional RNA molecules at least once during the experimental time, and two S7 binding-incompetent classes S7-/probe+ (with probe binding) and S7-/probe-(no probe binding), which we assign as misfolded and thus non-functional RNA molecules during the entire experimental time (Fig. 3D,E,F and fig. S6 and S7).

Under our experimental conditions, ∼50% of the molecules could bind S7 - similar for both the tested probes targeting H2829 (∼49%) and H30 (∼48%) regions on the rRNA, which is in agreement with our probes not affecting RNA folding. For the H30 region, only a minority (∼18%) of the S7 binding-competent RNA molecules also show probe binding, in agreement with successful H30 formation required for S7 to bind. In contrast, for the H2829 site, we see that ∼75% of the S7 binding-competent molecules have the H2829 site accessibility during the experimental time (Fig. 3F). This finding was unexpected, considering that H2829 is at the center of the S7 binding site and includes several intermolecular interactions between H2829 and S7 (Fig. 3B and fig. S5A). While we do not see concomitant S7 and H2829 probe binding, this major class of RNA molecules transitions back and forth between S7 binding competent and H2829 accessible states multiple times during the 20 minutes experimental time, suggesting a dynamic equilibrium between a folded and slightly unfolded S7 binding site at this central region during this early stage of ribosome assembly (fig. S6). Mutations introduced to stabilize H2829 caused reduced fraction of molecule with H2829 accessibility, while mutations that decreased H2829 stability decreased the fraction of S7 binding molecules, effectively shifting the equilibrium (fig. S8).

Our data allow us not only to assign different RNA conformations into different nascent RNA folding classes but they also provide information on the progression of nascent RNA maturation from co-transcriptional RNA folding to post-transcriptional RNA relaxation following dissociation of the nascent RNA from the transcription machinery. We plotted the aggregated probe accessibility over time for the RNA folding class S7+/H30+ (see methods). We find that in the first 50 s, which coincides with the median time that the nascent RNA is associated with the transcription machinery, there is substantially increased H30 accessibility till it subsequently decreases as soon as S7 starts to bind (Fig. 3 I,J and fig. S7 and S9A). In contrast, for the S7+/H2829+ class, we do not detect increased probe binding to early nascent RNA but rather see probe binding during the entire experimental time (fig. S6 and S9C). This is in agreement with a dynamic interchange between two conformations, one with high H2829 linker accessibility and one with S7 binding, as described above.

### Secondary r-proteins guide folding by increasing local RNA accessibility

In order to understand how the RNA folding classes change during the course of ribosome assembly, we next investigated the effect of secondary r-proteins (S9, S13, S19) on the co-transcriptional rRNA folding dynamics. Ribosome assembly is a sequential process in which secondary r-proteins, such as S9, S13, S19, can only detectably bind after successful binding of primary ones, such as S7 (*31, 39, 51*). As expected, the fraction of S7-binding competent molecules increased from ∼50% to ∼80% in agreement with an RNA chaperoning function of the secondary r-proteins for the 3’domain (Fig. 3G,H and fig. S9D,E) (*25*). Furthermore, the median *K*_d_ of S7 for the individual RNA molecules decreased for both S7+/probe- and S7+/probe+ classes by 1.5 fold/1.8 fold (for H2829 site) and 1.8 fold/1.2 fold (for H30 site), respectively (fig. S10A-D). In agreement, the median S7 arrival times per molecule decreased (fig. S10E,F) and S7 bound-lifetimes (fig. S10G,H) as well as longest S7 bound dwell time per RNA molecule (fig. S11) increased in presence of the secondary binding r-proteins, to a similar extent for all S7+ folding classes (S7+/probe- and S7+/probe+).

Apart from the expected increase in the fraction of S7-binding competent RNA molecules, unexpectedly, we also observed a substantial increase (∼20% to ∼36% of total molecules) for the fraction of molecules accessible at H30 upon addition of secondary r-proteins, while accessibility to H2829 stays high (∼73% to ∼69% of total molecules). By separately analyzing the four RNA folding classes, we find that mainly the subset of RNA molecules with accessible H30 is responsible for the increased fraction of S7 binding competent RNA molecules in presence of the secondary r-proteins (S7+/H30+ class: 3.6 fold increase; S7+/H30-class: 1.3 fold increase), while the fraction of S7-binding competent molecules increases independently of H2829 accessibility (both S7+/H2829- and S7+/H2829+: 1.5 fold increase) (Fig. 3G,H and fig. S9D,E). Since H30 is surrounded by the binding sites of the secondary r-proteins (fig. S1A), we hypothesize that their transient binding could rescue stable H30-dependent RNA folding traps (see supplementary text) (*52, 53*), thereby making the region around H30 more dynamic as experimentally detected by more transient DNA probe binding to this region. Overall, simultaneously tracking site-specific RNA accessibility and functionality of the same RNA molecules, we find how secondary r-proteins change the nascent RNA conformational landscape distribution, whereby increased conformational dynamics at specific sites correlate with increasing rRNA folding efficiency mediated by the secondary r-proteins.

### ASOs and assembly factors can specifically modulate a subset of RNA folding classes

Next, we tested if the different classes of RNA molecules can specifically be modulated by artificial as well as endogenous ligands. We first investigated the effect of ASOs binding to the upper part of the 3’domain (intervening RNA flanking H30; Fig. 4A). This region lies outside of the S7 binding site and was shown not to be required for S7 binding, but when this region was present, it decreased the co-transcriptional rRNA folding efficiency by promoting rRNA misfolding (*25*). We therefore hypothesized that ASOs binding to this region would increase the fraction of S7 binding competent molecules by sequestering this upper region thereby reducing co-transcriptional rRNA misfolding of the extended S7 binding site (Fig. 4A). Unexpectedly, we find the opposite: A single ASO binding to H31 reduced the fraction of S7 binding competent RNA molecules 2.1-fold (∼48% to ∼23% decrease for total RNA molecules of combined S7+/H30- and S7+/H30+ classes). Notably, the effect is class-specific: while the S7+/H30-class reduces substantially (5.6-fold decrease: ∼40% to ∼7%) there is, in contrast an increase for the S7+/H30+ class upon ASO addition (1.8-fold increase: ∼9% to ∼16%) (Fig. 4B,C and fig. S12A,B). Also, ASOs binding to other sites differentially affect the RNA folding classes, overall illustrating the complex redistribution of the nascent RNA conformational ensemble upon site-specific ASO binding.

**Fig. 4:**
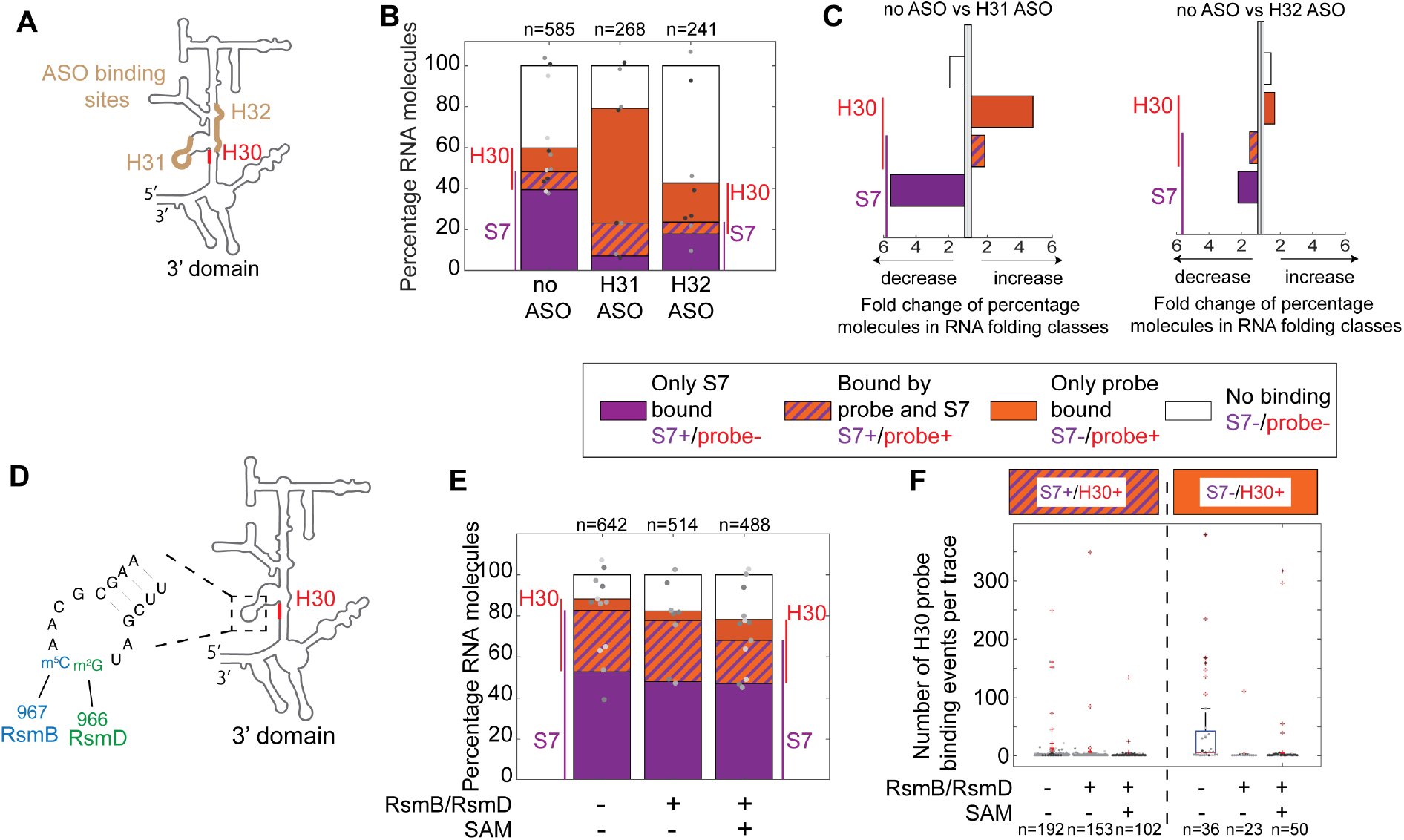
Modulating different RNA folding classes by assembly factors and ASOs. **(A)** Schematic of the 3’domain with H30 site being probed (red) and the ASO target sites (brown). **(B**,**C)** Percentage of molecules in RNA folding classes **(B)** and relative fold change **(C)** in presence and absence of different individual ASOs. **(D)** Schematic of the 3’domain with H30 site (red) and the corresponding nucleotides modified by RsmB (blue) and RsmD (green). **(E)** Percentage of molecules in RNA folding classes in absence or presence of RsmB/RsmD and/or SAM (always in presence of S9, S13 and S19). **(F)** Boxplot of number of H30 binding events per trace for S7+/H30+ (orange and purple stripes) and S7-/H30+ (orange) folding class in presence of secondary r-proteins with different combinations of RsmB/D and/or SAM. **(B**,**E**,**F)** Number of molecules analyzed (n) is shown. Bars are calculated by pooling of 3,2,2 **(B**,**C)** and 3,2,3 **(E**,**F)** replicates respectively, shown as tones of grey dots. No ASO and no RsmB/RsmD/SAM condition is the same as presented in Fig. 3G.

Next, we wondered whether ribosome assembly factors that bind in the vicinity of the functionally important H30/H31 region could affect co-transcriptional rRNA folding. We chose the rRNA modification enzymes RsmB (methylating 967C to 967-m^5^C) and RsmD (methylating 966G to 966-m_2_G) presumably acting during early stages of 3’domain assembly (Fig. 4D) (*54, 55*). While they minimally affect the relative fraction of our 4 defined RNA folding classes (Fig. 4E and fig. S12C,D,F,G), they specifically and substantially reduce the number of H30 probe binding events (change of median number of binding events per trace: 5.5 to 1; and 75-percentile: 42.5 to 2 in absence and presence of RsmB/D, respectively; Fig. 4F) and showed a 3.2-fold reduction in the accessibility score (0.049 to 0.015; fig. S12E) for the S7-/H30+ class, an effect which is independent of the presence of the co-factor SAM required for the methylation (Fig. 4F and fig. S12C-E). We hypothesized that by transiently sequestering the loop of H31, RsmB/D prevent the formation of the misfolded S7-/H30+ subclass with high probe accessibility. To test this hypothesis, we perturbed the RsmB/D binding site by replacing the H31 loop by an UUCG tetra-loop (fig. S13A). This minimally affected the RNA structural ensemble but reduced again the number of H30 probe binding events in molecules of the S7-

/H30+ class substantially, being phenotypically equivalent to the presence of RsmB/D (fig. S13B,C). These findings support a model in which loop H31 promotes the formation of a specific misfolded RNA class if not transiently sequestered by assembly factors or mutated. The effect of RsmB/D on only a small subset of the RNA molecules would be undetectable in bulk experiments, thus, providing a first biophysical rational for the minimal growth phenotype under normal growth conditions for a RsmB/D double deletion mutant (*56*). Overall, these data demonstrate how various nascent rRNA folding classes can specifically be modulated by artificial or natural ligands.

### Dynamic multi-site RNA structure probing with functional readout

We showed that different RNA sites have different tendencies to remain dynamically accessible in functional RNA molecules. However, it is not clear whether different sites cooperate to be transiently accessible. In order to simultaneously track the dynamic accessibility of two sites and in addition, assign the functional state of the RNA molecules, we repeated our co-transcriptional RNA probing experiments by starting the reaction with the addition of NTPs, Cy7-H2829-DNA probe, Cy5-H30-DNA probe and Cy5.5-S7 r-protein, requiring the simultaneous detection of 5 colors and the acquisition of >400 molecules for 30 minutes (Fig. 5A,B and fig. S14A,B). Analogous to our above presented experiments, we classified the different RNA molecules into all possible accessibility and functional states, now resulting into 8 possible classes: +/-H2829 accessible, +/-H30 accessible, +/-S7 binding-competent and combinations of these (fig. S14C top rasterplot). We found molecules in all the 8 classes, with the aggregated fraction of molecules in agreement with experiments in which only a single site was probed together with S7 functional readout (fig. S15A), using different labeling schemes (fig. S15B,D) and for different replicates (fig. S15C), thereby validating our complex multicolor datasets and further confirming that various combinations of DNA probes used here do not detectably affect RNA folding dynamics.

**Fig. 5:**
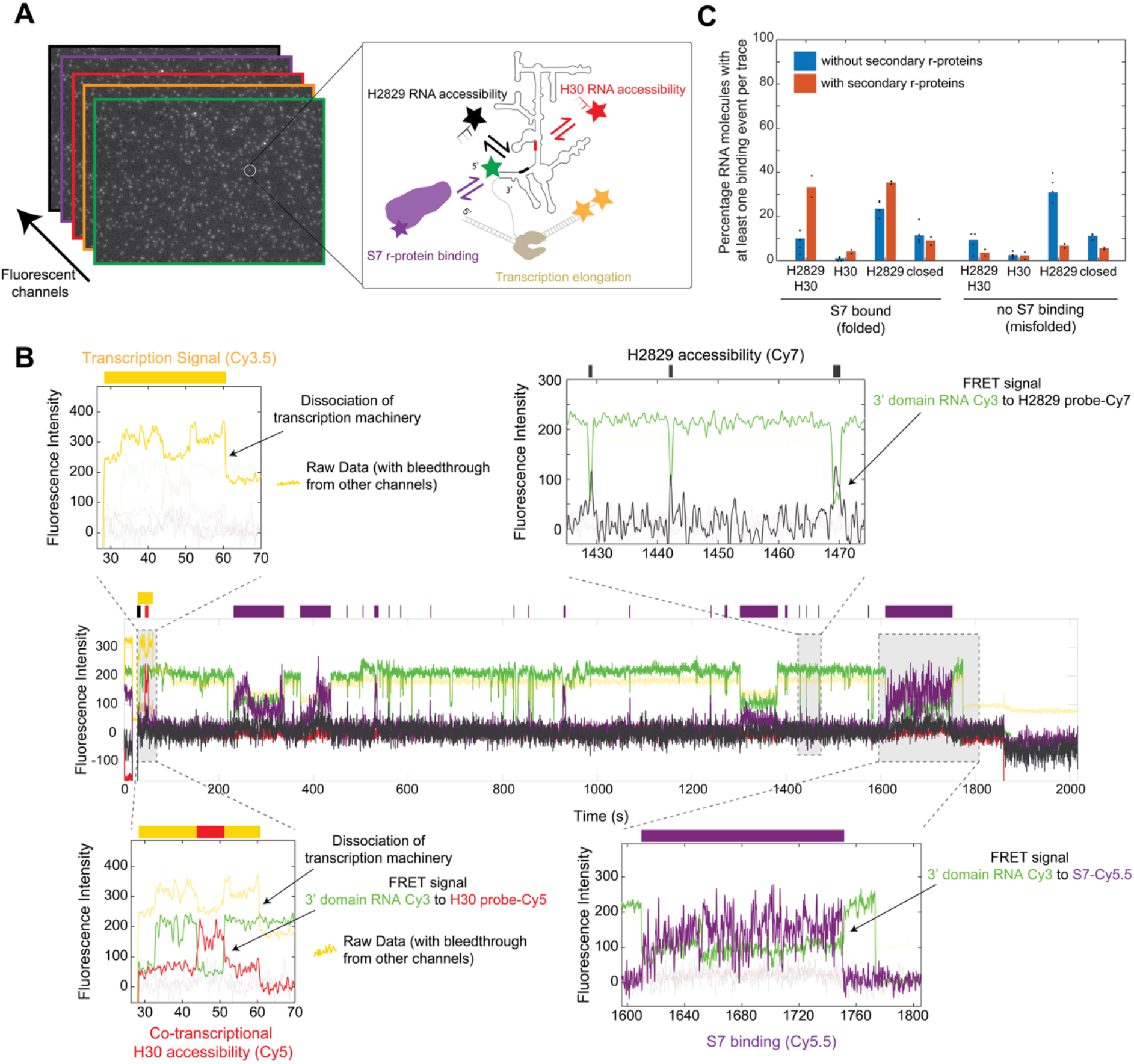
Multi-site nascent RNA probing with functional readout. **(A)** Experimental setup for simultaneously probing transcription elongation, H2829 and H30 RNA site accessibilities and the binding of r-protein S7, acquired from 5 fluorescent channels. **(B)** An example smoothed trace of the 5-color data showing FRET donor (green – Cy3 dye), transcription signal (yellow – Cy3.5 dye), H30 probe (red – Cy5 dye), S7 r-protein (purple – Cy5.5 dye) and binding of the H2829 probe (black – Cy7 dye). Simplified representation of binding events is shown on top of the trace as bars. Insets zoom into specific regions highlighting the specific signals. **(C)** Percentage of RNA molecules within each of the 8 folding classes in presence and absence of secondary r-proteins. Plotted by pooling 4 replicates – without, 2 replicates – with secondary r-proteins.

Our data show that H2829 can be accessible independent of H30 accessibility, both for S7 binding competent and binding incompetent molecules (Fig. 5C, fig. S14C bottom rasterplot). Repeating our experiments in presence of secondary r-proteins supports our finding that different sites can be accessible independently. We find that S7 binding competent classes which are accessible by either H30 and/or H2829 probes increase in abundance in presence of secondary r-proteins, while the abundance of the S7 binding competent class which is inaccessible to both DNA probes does not substantially change in presence of secondary r-proteins (Fig. 5C, fig. S14C). Overall, these findings further support a model in which transient accessibilities at local RNA sites correlates with the ability of secondary r-proteins to chaperone nascent rRNA folding.

## Discussion

Using up to 5-color single-molecule fluorescence imaging, we describe part of a dynamic RNA conformational ensemble sampled as nascent rRNA emerges from the RNA polymerase and subsequently continues folding post-transcriptionally. Our data show how subsets of RNA conformations can differentially be modulated by various factors (Fig. 6). Unexpectedly, we find that RNA molecules with higher local dynamic accessibilities, constituting transient non-native structures, have a higher propensity to be chaperoned by secondary r-proteins and thus, are overall more efficiently folded. In vivo, RNA is less structured and more dynamic, likely due to transient binding of RNA binding proteins and ATP-dependent helicases (*57*). Overall, our data is consistent with a model in which dynamic RNA structure provides an advantage to form productive RNPs by minimizing RNA folding traps.

**Fig. 6:**
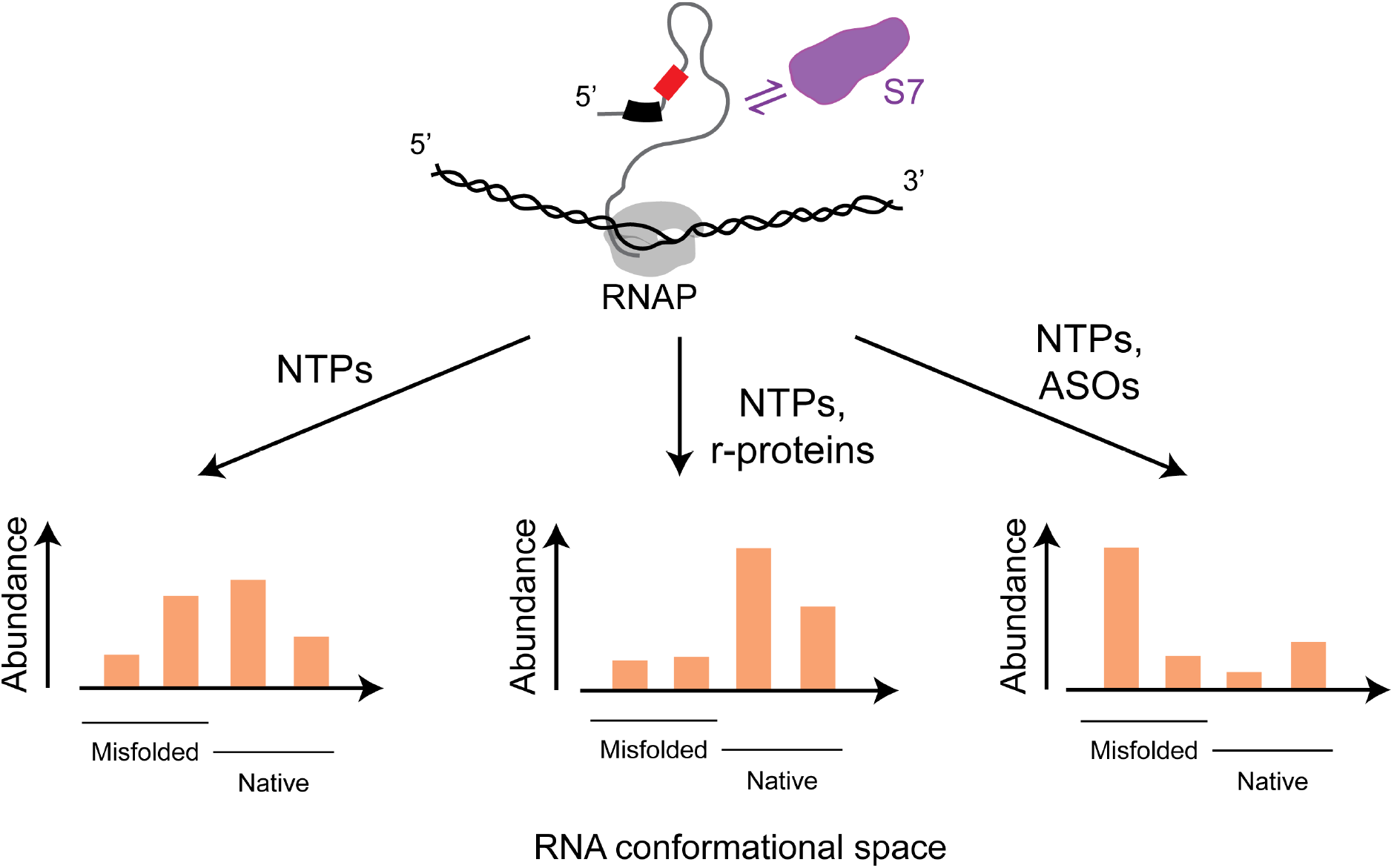
Ligands shape the nascent RNA conformational distribution landscape. The simplified model shows the effect of various factors on the distribution of the rRNA conformational landscape.

Previous biochemical and structural studies have investigated the heterogeneity in the rRNA folding and r-protein binding landscape using pre-transcribed and pre-folded RNA (*40-44, 49, 58*). Few studies have directly visualized the heterogeneity in r-protein binding to the nascently transcribed rRNA (*24, 25, 28*). Our experiments systematically dissect the co-transcriptional and post-transcriptional RNA structural heterogeneity at various levels. First, comparing individual RNA molecules of the structural ensemble, we show that different nascent RNA molecules probed at an individual site have high variability (>3 orders of magnitude) in their accessibility profiles (Fig. 2C). Second, comparing RNA accessibility across different sites further illustrates the heterogeneous behavior of the RNA. Third, simultaneously tracking rRNA accessibility and r-protein binding on the same nascent rRNA molecules revealed the presence of different RNA folding classes that would have been missed by only tracking RNA structure or protein binding at a time. Fourth, the molecules show heterogeneity in RNA folding and r-protein binding kinetics across molecules within a class as well as across different classes (Fig. 3). Finally, simultaneously tracking the accessibility of two RNA sites with additional functional readout, we classified our dynamic structural ensemble into 8 dynamic RNA folding classes, differing in both structural (local site accessibilities) as well as functional properties (the ability for a protein to bind the target) of single RNA molecules, which further highlights the nascent RNA structural heterogeneity (Fig. 5). However, the true structural diversity is much higher as we only detect information on a subset of all the sites at the same time. This structural diversity makes specific targeting of RNA structure much more challenging. Initial attempts to target RNA with small molecules were successful so far only for locally rigid structures such as GUC-repeats, the 5’splice-site of SMN2 exon 7 or the ribosome, but more dynamic systems such as nascent rRNA present during ribosome assembly still await successful targeting (*59-62*). Successful targeting of RNA will require an ensemble description of RNA structure, as suggested by improved targeting of local RNA structural ensembles instead of a single static structure (*63-66*).

More promising is targeting of RNA with anti-sense oligonucleotides (*67*). We found a high variation in the fraction of RNA molecules that were accessible to our short hybridization probes at different local sites on the RNA. These findings provide a direct biophysical visualization and explanation of why certain sites in long mRNA molecules are more targetable by ASO than other sites (*68-74*).

While our data only allows us to classify the heterogenous pool of nascent RNA conformations into specific folding classes based on few local RNA accessibilities and the binding of S7, we can compare our dynamic and functional data to previous static RNA structure probing data. We found that the H30 probe binding is highly enriched during transcription, but binding is reduced as soon as S7 can bind (Fig. 3I,J). Furthermore, destabilizing the region surrounding H30 resulted in a 5.6-fold reduction in S7 binding (Fig. 4 and fig. S13). Both findings suggest that H30 needs to be formed for S7 to bind. This is in agreement with footprinting data showing that H30 exhibits increased protection upon S7 binding despite lacking direct interactions between S7 and H30 (*34*), thus linking H30 formation with S7 binding.

Furthermore, hydroxyl radical footprinting of isolated native 16S rRNA in the presence of all 30S r-proteins showed that S7 forms immediate interactions with H43 but engages more than an order of magnitude slower with the H2829 junction (*40*). These data, together with our observations that post-transcriptionally, H2829 accessibility dynamically interchanges with S7 binding points to a model in which the H2829 junction is not stably pre-formed for S7 to engage immediately, but rather can switch between various open and closed conformations. These observations are also in agreement with S7 binding relatively late during 30S assembly despite being the primary binding protein of the 3’domain (*75, 76*).

In addition, we demonstrate that a single H31 ASO binding outside the S7 binding site strongly inhibited S7 binding and simultaneously increased accessibility of a neighboring RNA site (H30) in this misfolded (S7 binding incompetent) class of RNA molecules (6-fold decrease in S7+/H30- and 5-fold increase in S7-/H30+ class), while no RNA folding class changed by >2-fold for the H32 ASO (Fig. 4C). Interestingly, footprinting data show that nucleotides bound by the H31 ASO strongly change reactivities upon addition of S7 and further upon addition of secondary r-protein S19 (*34*). Furthermore, H31 ASO encompassing nucleotides fold more slowly than most nucleotides on the H32 ASO binding site as measured by hydroxyl radical footprinting of the entire 16S rRNA in the presence of all 30S r-proteins (*40*). Finally, H31 forms part of the core of the 3’domain and requires assembly factor RimM for proper folding in vivo (*77*), highlighting its sensitivity to perturbations. Overall, these observations suggest that targeting H31 or the surrounding regions by an ASO has a large effect on RNA folding, as this helix is a critical checkpoint in 3’domain folding.

Our structural and dynamic characterization of molecules into various RNA folding classes not only enables us to describe native (S7 binding competent) RNA molecules, but also provides information on the pool of misfolded (S7 binding incompetent) RNA molecules. Research has focused on elucidating the functional native structures, because these are physiologically relevant and easier to study as they are usually present in a single well-defined conformation. In contrast, very little is known on the structural diversity and properties of misfolded RNA structures ((*25, 28, 78-80*) and this work). Here, we also investigate the pool of RNA conformations that misfold during nascent RNA folding. For example, we observe a small class of molecules (S7-/H30+) that are kinetically trapped as they are accessible at H30 over the entire duration of the experiment and never refold into S7 binding competent conformations, irrespective of the presence of the r-protein chaperones (Fig. S9B). Kinetic traps have been described for large RNAs like for the Tetrahymena ribozyme where incorrect topology leads to a long-lived misfolded state with the critical catalytic domain being mispositioned requiring extensive refolding for the misfolding to be resolved (*78, 79, 81-83*). The inability of some nascent rRNA molecules to rearrange over 30 minutes indicates that the RNA is highly structured, which may prevent “free ends” to initiate strand displacement, a mechanism for efficient rescue of RNA folding traps (*53, 84-86*).

In contrast, the molecules in the S7+/H30+ class are initially accessible at H30 and then change to a S7 binding competent conformation. Furthermore, this effect is enhanced in the presence of r-protein chaperones. We associate this with the ‘pre-association’ chaperoning mechanism of the r-proteins, where the transient association of the protein caused by non-specific or a subset of specific interactions prevents the RNA from forming stable non-native interactions as a mechanism to rescue formation of folding traps (*52*). These weak non-native interactions could efficiently be rearranged via RNA strand displacement (*53*). We also observe high variation in accessibility across different sites. For example, unlike H30, the molecules of S7+/H2829+ class keep transitioning between a partially misfolded H2829 site and a S7 binding competent state. This local misfolding occurs at shorter timescales in contrast to the above-described stable kinetic RNA folding traps.

Overall, understanding mechanisms on how RNA can specifically misfold holds great promise for novel future strategies to target RNA: next-generation drugs could be designed to steer RNAs into, and specifically stabilize these misfolded and non-functional conformations to modulate cellular function.

## Acknowledgments

We thank Martyn Reynolds and Felix Jan Evers from Cairn Research/ Ultimeyes for the collaborative effort in setting up the multi-color TIRF microscope; Marko Lampe from the EMBL Advanced Light Microscopy Facility, EMBL IT team, Thomas Hoffmann, Andrey Revyakin, the EMBL chemical core facility for input or assistance in setting up microscope pipelines. We also thank the EMBL protein expression and purification core facility, Benjamin Lau and Nusrat Qureshi for purification of recombinant *E. coli* RNA polymerase and r-protein S10 and S7, respectively. We thank the entire Duss lab for helpful discussions. We also thank Janosch Hennig and Zeynep Baharoglu for critically reading the manuscript.

## Author contributions

Conceptualization: OD Methodology: OD and KG Investigation: KG Visualization: OD and KG Funding acquisition: OD Software: EMG, KG, OD Supervision: OD Writing – original draft: KG, OD Writing – review & editing: OD, KG

## Data and materials availability

All data are available in the main text or the supplementary materials. Scripts for downstream processing of single-molecule traces were made available publicly previously under an open-source license (*87*). Software for single-molecule data processing is published and freely available online (*88*). The large, raw video files are available on request from the corresponding author. Structures were downloaded from the Protein Data Bank (http://www.rcsb.org/) using the accession code show in Fig. 3B, fig. S1 and S5 (4V9P).

