## Supplementary Material for "Functional characterization of dynamic nascent RNA folding ensembles in real-time"

Science Advances
AAAS

### Supplementary Materials for

#### Functional characterization of dynamic nascent RNA folding ensembles in real-time

Kavan Gor[1,2], Eva Maria Geissen[3], Olivier Duss[1,2]*

**The PDF file includes:**

Materials and Methods
Supplementary Text
Figs. S1 to S15
Tables S1 and S2
References 1 to 16

#### Materials and Methods

Experimental model

Plasmids for *E. coli* r-proteins S3 (WT and M129C), S7 (WT and S83C with a truncated C terminus (157-178)), S9, S10, S13 and S14 were a kind gift from the Williamson lab (*1*). The genes for *E. coli* S19, RsmB and RsmD were cloned into a pESUMO vector backbone using Gibson assembly. Gene sequences were obtained from the ASKA collection plasmids (*2*).

DNA templates and in vitro transcription

The DNA transcription templates were designed as described previously (*3*). In brief, the 3' end of the DNA template was designed to have a single stranded overhang where a DNA oligo could be hybridized. The single stranded overhang was generated using autosticky PCR using a gene fragment ordered from IDT (p0030_ab_fw and p0075_ab_bw). The PCR primers were designed to have a complementary sequence to the DNA template, an abasic site and the complementary sequence for hybridization to a labelled DNA oligo. PCR was performed using Phusion DNA polymerase as described in the manufacturer's instructions. The presence of a single PCR product was verified on an 1% agarose gel, which was then purified using a Qiagen PCR purification kit following the instructions provided. If the PCR reaction contained side products, the correct product was purified from a 1% agarose gel using a Qiagen gel extraction kit, followed by buffer exchange into 10 mM TrisHCl (pH 7.5) and 20 mM KCl using a 30 kDa molecular weight cutoff centrifugal filter (Amicon).

A 1.2x excess of DNA oligo labelled with two Cy3.5 fluorescently dyes (p0088_2xCy3.5, ordered from IDT) was used to hybridize to the single-stranded overhang of the DNA template by annealing at 68 °C for 5 minutes and slow cooling at room temperature.

Protein expression and purification

The proteins were purified in denaturing conditions as described earlier (*1*). In short, the proteins (S3, S7, S9, S10, S13, S14) were expressed in *E. coli* (BL21(DE3) Gold strain) at 37 °C in LB medium and the cells lysed in a microfluidizer. The resultant inclusion bodies were washed, then solubilized in 6M guanidine chloride or 8M Urea (for r-protein S9 and S10) and dialyzed overnight in 8M urea. The purification was then continued in denaturing conditions where samples were loaded on a SP HiTrap column (GE Healthcare) or a Ni-NTA column (for r-proteins S9 and S10) followed by a SP column (for S9 r-protein). r-protein S10 was dialyzed overnight to cleave SUMO tag at 4 °C, followed by reverse Ni-NTA chromatography. The r-protein S10 was concentrated and dialyzed in the final buffer (50 mM TrisHCl (pH 7.5), 400 mM KCl, 2 M Urea, 7 mM betamercaptoethanol) and frozen at -80 °C. The S3, S7, S9, S13, S14 r-proteins were dialyzed overnight in refolding buffer (20 mM TrisHCl (pH 7.6), 20 mM NaCl, 0.5 mM EDTA, 0.5 mM DTT) at 4 °C. The folded protein was then purified using a Heparin HP column (for r-protein S3, S7 and S14) followed by size-exclusion chromatography on a Superdex 75 26/600 HiLoad gel filtration column (GE Healthcare) equilibrated with final sample buffer (20 mM TrisHCl (pH 7.6), 100 mM NaCl, 0.5 mM EDTA, 0.5 mM DTT). r-protein S19, as well as RsmB and RsmD were expressed in LB medium at 37 °C, cells lysed and the lysate loaded on a Ni-NTA column (Cytiva), followed by overnight dialysis at 4 °C to cleave the SUMO tag and reverse Ni-NTA chromatography. Proteins were further purified using a SP column (for r-proteins S19) or using size-exclusion chromatography on a Superdex 75 26/600 HiLoad gel filtration column (GE Healthcare) equilibrated with final sample buffer (50 mM TrisHCl (pH 8), 200 mM NaCl, 0.5 mM

EDTA, 0.5 mM DTT) (for RsmB). The monomeric fractions were pooled and concentrated using an Amicon 3-10 kDa MWCO Ultra Centrifugal Filter (Millipore) depending on the size of the protein and frozen at -80 °C.

Protein fluorescent-dye labelling

The S7(S83C) variant with a truncated C terminus (157-178) and S3(M129C) variant were labelled using Cy5.5 sulfo-maleimide in denaturing conditions as previously described (*1*). Briefly, 1 mg of r-protein was reduced using 10 mM DTT on ice for 2h in labelling buffer (100 mM $Na_2HPO_4/KH_2PO_4$ phosphate buffer pH 7.0, 100 mM NaCl), followed by ammonium sulphate precipitation (70 % w/v) and incubation on a rocker for 20 minutes on ice. The protein was pelleted (14000 rpm for 20 minutes) and washed with 70 % ammonium sulphate in labelling buffer and again centrifuged at 14000 rpm. The washed pellet was resuspended in labelling buffer containing 6M urea. r-protein was mixed with 1 mg of Cy5.5 sulfo-maleimide and incubated on a shaker (50 rpm) at room temperature for 30 minutes and the reaction stopped by adding 0.5 % betamercaptoethanol. Excess dye was removed by purifying the reaction on a Nap5 column (GE Healthcare) equilibrated with labelling buffer containing 6M urea. The resultant fluorescent fractions were visualized on a 4 %-20 % SDS-PAGE using a Typhoon imager (Cytiva) and with Coomassie stain. The r-protein was refolded by diluting the sample 10-fold in sample buffer (20 mM TrisHCl (pH 7.6), 20 mM NaCl, 0.5 mM EDTA). A second purification step was performed by passing the sample through a Heparin HP column at 4 °C. The column was washed with 10 column volumes of sample buffer and the protein was eluted with 1 M NaCl in sample buffer. The fractions were visualized as mentioned above. The labelled fractions were pooled, aliquoted and stored at -80 °C.

Preparation of stalled Transcription Elongation Complex (TEC) for single-molecule imaging

The TEC for single-molecule imaging was generated as described earlier (*3*). To summarize, a paused transcription elongation complex consisting of a DNA template, a single RNA polymerase and a nascent RNA of 50 nucleotides (containing adapter sequences) was generated by incubating 25 nM DNA transcription template, 100 nM *E. coli* RNA polymerase (in-house generated or purchased from NEB), 100 μM ACU trinucleotide (Dharmacon), 10 μM ATP, CTP and UTP (Jena biosciences), 2 mM DTT in a buffer containing 50 mM TrisHCl (pH 8.0), 14 mM $MgCl_2$, 20 mM NaCl, 0.04 mM EDTA, 40 μg/ml BSA (non-acylated), 0.01 % Triton X-100 for 20 minutes at 37 °C (references(*3, 4*)). Then, 20 nM of a pre-annealed dsDNA oligo construct containing a biotin at the 5' end and a single-stranded overhang at the 3'-end (p44 and p109-biotin) complementary to the adapter region on the nascent RNA was added to be annealed to the paused elongation complex along with 1 mg/ml heparin to prevent transcription re-initiation. The mixture was incubated at 37 °C for 20 minutes.

In order to perform the single molecule assay, the microfluidic channel was functionalized and assembled as described earlier with a small adaption (*5*). Each channel was prepared such that it consisted of two inlets and two outlets. The channel and the inlet and outlet tubing were first washed with buffer (10 mM TrisHCl (pH 8.0), 50 mM NaCl) and incubated with 0.1 mg/ml Neutravidin for 5 minutes, followed by incubating the TEC for 10 minutes in imaging buffer (50 mM TrisHCl (pH 7.5), 14 mM $MgCl_2$, 20 mM NaCl, 0.04 mM EDTA, 40 μg/ml BSA (non-acylated), 0.01% Triton X-100, 2 mM spermidine, 1 mM putrescine and 150 mM KCl).

Probe design

The probes were designed such that they have only one site that is fully complementary (containing 7 consecutive nucleotides) and do not have alternate target sites with more than 4 consecutive nucleotides complementary in the entire 3'domain to minimize the detection of off-target probe binding (fig. S2).

<u>Single-molecule co-transcriptional RNA accessibility assay</u>

In order to track DNA probe and protein binding during and following transcription, we had to hybridize a Cy3-DNA oligonucleotide to the 5'-end of the nascent RNA: After immobilizing the stalled TEC (ATP, CTP, UTP stalled) to the imaging surface and washing away excess nucleotides, we walked the RNAP (10 minutes at 21 °C in imaging buffer containing 10 µM ATP, CTP, GTP and 100 nM p66-Cy3-FQ oligo or p153-Cy3B-FQ only when specified) such that the Cy3-oligo binding site on the nascent RNA becomes accessible for hybridization by the oligo. Finally, the slide was washed with imaging buffer containing oxygen scavenging mix (OSC) consisting of 2.5 mM protocatechuic acid (PCA) and 250 mM protocatechuate-3,4-dioxygenase (PCD) to prevent photobleaching and 4 mM Trolox as triplet state quencher, ready for single-molecule imaging (*6, 7*).

The assembled slide with immobilized molecules was used to focus molecules on the custom-built multi-color Total Internal Reflection Fluorescence (TIRF) microscope. The parameters were setup (see instrumentation section) such that a 10 seconds laser dead time was encoded after 15 seconds from the start of the acquisition. The laser dead time was used to inject the reaction mixture (to prevent photobleaching of the immobilized molecules) containing 1 mM NTPs to re-initiate transcription elongation, 200 nM (or specified concentrations) of each of the labelled short DNA probes (to probe RNA accessibility, H2829 probe: p110Bcy5 or prKG058cy5.5 or prKG080cy7; H30 probe: prKG032cy5 or prKG057cy7; H32 probe: prKG053cy5) and/or 20nM S7-Cy5.5 (to provide functional readout), and/or 400 nM of each of the unlabeled secondary r-proteins, and/or 400 nM of each of the ASOs (H31: prKG074; H32: prKG073; H34: prKG072; H30-H41 junction: prKG075; H42-H29-H43 junction: prKG076; H28: prKG077), and/or 400 nM of each of the unlabeled RsmB and RsmD proteins. The chase mix was prepared in the imaging buffer containing 0.25 % Biolipidure 203, 0.25 % Biolipidure 206, 0.5 mg/ml yeast total tRNA and OSC (see above). In order to distinguish single molecules, as compared to multiple molecules present on a single diffraction limited spot, another reaction mixture was injected in the last 1 % of the acquisition time. The second mixture was made in imaging buffer but containing only 0.25 mM PCA, 25 mM PCD and 0.4 mM Trolox.

<u>Single-molecule assay to probe single-stranded RNA/DNA (ssRNA/DNA) accessibility</u>

Single-stranded RNA/DNA labelled with Cy3 on one end and biotin on the other end and containing the region of interest of the 3'domain was immobilized (ssRNA of H2829: p112A; H30: prKG047; H32: prKG078; ssDNA: prKG081). The reaction mixture made in imaging buffer consisted of 0.25 % Biolipidure 203, 0.25 % Biolipidure 206 and 0.5 mg/ml yeast total tRNA (except for experiments in fig. S1B); 200 nM of the labelled probe and OSC mix were injected after the start of acquisition. The second injection mixture made in imaging buffer but containing only 0.25 mM PCA, 25 mM PCD and 0.4 mM Trolox was injected in the last 1 % of the acquisition time to induce photobleaching.

<u>Single-molecule assay to probe RNA accessibility in the pre-assembled 3' domain</u>

To assemble the 3' domain, 400 nM of unlabeled r-proteins S7, S9, S13, S19, S10, S14 and 400 nM of labeled S3(M129C)-Cy5.5 was added to the 3' domain transcription reaction or to the 3'domain rRNA that was pre-transcribed and pre-folded at 37 °C. The reaction was incubated for 30 minutes. The pre-assembled complex was immobilized and used for probing RNA accessibility at 21 °C as described above. Successful assembly of the 3'domain was identified by S3(M129C)-Cy5.5 bound at the beginning of the experiment (Cy3-Cy5.5 FRET) as described previously (*1*).

Instrumentation and data acquisition parameters

All the single-molecule experiments were acquired on a custom-built objective type (CFI SR HP Apochromat TIRF 100XC Oil) TIRF on an ECLIPSE Ti2-E inverted microscope (Nikon), built in collaboration with Cairn Research: https://cairn-research.co.uk/ and Ultimeyes: https://www.ultimeyes.eu/. Homogenous illumination over the full field of view and modulation of laser power was achieved using an iLAS modular scanning system interface (GATACA systems). The total fluorescent signal was split into their respective fluorescent channels based on the dye emission spectrum using a combination of bandpass and emission filters and the photons detected using three Prime95B sCMOS cameras (Teledyne Photometrics) with an exposure time of 100 ms at 21 °C. All the experiments were acquired using an OBIS 532 nm LS 150 mW laser head with an output intensity of 0.2 kW $cm^{-2}$ at the objective.

Single-molecule data analysis

Single-molecule data from different cameras (two colors per camera) were split into stacks of individual fluorescent channels using the Metamorph software. Subsequently, the different fluorescent channels were organized into a single tile. The stacks were saved as Metamorph Multitiff files. The tile stacks were loaded into a modified version of the SPARTAN software (*8*) implemented in MATLAB. The localized molecules were visually inspected to check for drift over the duration of the experiment and drift corrected where necessary using an in-house written python script. The resultant drift-corrected files were processed with SPARTAN to perform channel alignment and registration as well as extraction of the individual single molecule trajectories. The single molecule trajectories were then further analyzed using MATLAB scripts as previously described (*1, 3, 9, 10*).

The molecules were selected based on specific criteria. For the co-transcriptional RNA accessibility experiments, the molecules were selected for two criteria: 1) molecules showing a characteristic gradual increase in fluorescent intensity of Cy3.5 (indication of transcription elongation) (*3*) followed by a single step drop in the fluorescent intensity (indicating DNA dissociation and thus dissociation of the transcription machinery) within 200 seconds. 2) molecules that have a Cy3 signal showing a single step photobleaching (indicating presence of only a single molecule at the diffraction limited spot) within the last 10-20 % of the total experimental time. For experiments where only RNA was immobilized, the traces were selected based on a single photobleaching step of the Cy3 oligo signal occurring during the last 10-20 % of the total experimental time.

Specific binding of dye-labelled short DNA probes (Cy5, Cy5.5 and Cy7) and/or dye-labelled protein (Cy5.5) was assigned using an anticorrelated high FRET signal between the Cy3-donor dye (Cy3-labelled oligo attached to the 5' end of the nascent RNA) and the acceptor dye (Cy5, Cy5.5 and/or Cy7). FRET efficiency was calculated as follows: $E_{FRET}=I_A/(I_A+I_D)$, where $I_A$ and $I_D$ are the apparent fluorescent intensities of the acceptor and donor dyes, respectively. Due to substantial spectral crosstalk between neighboring channels, the bound state was assigned using a threshold set at the midpoint of the two FRET states with subsequent manual inspection of traces that showed non-specific protein binding signal (causing a non-zero FRET efficiency baseline) as described previously in detail (*1, 3, 4*). In short, FRET efficiency was used as a guide to determine the specific binding, however for traces with non-specific protein binding, the anticorrelation of the individual donor and acceptor signal was used as a metric to distinguish the binding event from stickiness of protein to the surface resulting in high acceptor intensity but no simultaneous change in donor intensity.

Accessibility score

The accessibility score provides an indirect value for describing the relative accessibility of a single RNA site under various experimental conditions. We calculate the accessibility score the following: First, for each single trace we determine the fraction of the time the RNA molecule is bound by the probe. The accessibility score is the mean of this value averaged over all the molecules of an experiment or a specific RNA folding class.

Clustering

The rasterplots representing RNA accessibilities of individual sites in absence of r-proteins (Fig. 2D-F) were generated by hierarchically clustering molecules using Weighted Pair Group Method with Arithmetic Mean (WPGMA) and Spearman distance metric. The weighting of individual molecules was performed based on the number of short DNA probe binding events.

Time-dependent accessibility plots

After aligning the molecules post-experiment to the time of NTP delivery, the traces were binned into 5 seconds intervals and for each bin the percentage of molecules with 1) probe binding events and 2) S7 binding events were determined and plotted as a line plot. The median time from the start of transcription to DNA template dissociation for the molecules present in the specific S7+/H30+ class was extracted and overlayed on the plot (Fig. 3H,I).

Kinetic and thermodynamic analysis of S7 binding from single-molecule data

The on-rate determination of DNA or protein ligands binding to longer RNAs in single-molecule experiments is challenging due to incomplete labeling efficiency of the ligands and importantly, the presence of multiple RNA conformational states observed in individual RNA molecules. On-rate distributions are therefore best described by multi-exponential functions and thus challenging to interpret. Arrival times remain unaffected by these factors as they do not depend on any model. Thus, deriving $K_\mathrm{d}$ based on $k_\mathrm{off}/k_\mathrm{on}$ is challenging due to challenges in determining the on-rate. Therefore, we determine the apparent $K_\mathrm{d}$ from single RNA molecule binding to S7, assuming equilibrium of each RNA conformation, by using the fraction of bound RNA according to the below formula.

$$K_\mathrm{d,sm} = \frac{1-Y}{Y}\,[P] \qquad -(1)$$

$[P]$ describes the concentration of free protein. Y denotes the fraction of bound RNA ($Y = t_\mathrm{PR}/(t_\mathrm{PR}+t_\mathrm{R})$ with $t_\mathrm{PR}$ denoting the total time when RNA (R) is complexed with protein ($P$), and $t_\mathrm{R}$ denoting the total time when the RNA is free). Because the number of immobilized RNA molecules on the glass surface for single-molecule experiments is much lower than the total number of protein molecules added to the imaging solution, $[P] \approx [P]_0$ with $[P]_0$ being the total labeled concentration of the added protein in our single-molecule experiments. This was used to derive the $K_\mathrm{d}$ for individual molecules.

To determine the bulk $K_d$ across an entire class of RNA molecules, the molecules of the individual class were combined into a single molecule. The resultant bound time and unbound time were used to determine bulk $K_d$ using the above formula. The bulk $K_d$ was used to determine the fold change across different conditions. The data was evaluated between 200 s till donor photobleaching to calculate $t_{PR}$ and $t_R$ for both the above-mentioned $K_d$ analysis.

#### Supplementary Text

Comparing to existing single-molecule FRET approaches to monitor nascent RNA folding:

Previously (*1*), the formation of a long-range RNA helix, H28 in the 3'domain of the 16S rRNA, was tracked for a nascent RNA emerging from the RNAP. While this allowed determining which RNA molecules had H28 formed, it did not allow determining at what timepoint H28 formation happens, as explained in the following: Successful formation of long-range H28 was determined by detecting a Cy3/Cy3.5 FRET signal between Cy3 and Cy3.5 DNA oligonucleotides that were hybridized to artificial 19-22nts long sequences located at the 5'-end and 3'-end of the nascent RNA during transcription. However, the binding of the labelled DNA probes to the nascent RNA is very inefficient due to formation of secondary structure in the artificial probe binding region. This results into DNA oligonucleotide arrival times of dozens of seconds and thus, prevents determining the exact timepoints at which H28 forms and thus, does not allow for real-time tracking of RNA structure. Furthermore, this approach cannot be adapted for internal RNA structure but is limited to tracking the formation of long-range helices for which the RNA 5'-end and 3'-end base-pair.

Kinetic analysis of the probe binding to target sites:

In order to access the residence times of the probes, we fitted the bound dwell time distributions to single and double exponential functions. Comparing the fits of probe binding to the ssRNA and the 3'domain we observe that H30 and H32 show mostly single-exponential behavior as expected (fig. S1C). There is a slight deviation from perfect single-exponential behavior for H32 ssRNA, which is likely explained by some remaining RNA structure even in the small "ssRNA" construct, because probing of a ssDNA with identical sequence shows pure single-exponential behavior. H2829 probe binding dwell times to H2829 ssRNA follow a single-exponential kinetics; however, the dwell times to the 3' domain cannot be explained by a single-exponential fit. One explanation could be that for a fraction of the binding events, the H2829 target site is only partially accessible (5 or 6 out of 7 nucleotides) for probe binding, resulting in faster dissociation of the probe. Another possibility could be that the complementary RNA strand during local RNA folding could push away the DNA probe (reducing its bound lifetime) by strand displacement in agreement with our data showing dynamic opening and closing of the H2829 site (transitions between S7 and H2829). We also note that for sites H30 and H32, for which we see only a single exponential behavior, the off-rate is faster for the 3'domain compared to the isolated ssRNA constructs. We hypothesize that RNA conformational dynamics, such as local strand displacement, or incomplete accessibility of the probe for its 7 nucleotides binding site (i.e. only 6 nucleotides full complementarity), lead to generally faster DNA probe dissociation in the 3'domain RNA versus their isolated binding sites.

Our FRET-based probe detection approach detects all the probe binding events to the 3'domain rRNA:

rRNA folds and compacts instantly in presence of $Mg^{+2}$ (*11*) and therefore will never be fully unstructured and linearized, even if misfolded. The experimentally determined hydrodynamic radius of the entire 16S rRNA is 117 Å in 20 mM $Mg^{+2}$ measured at 25 °C (*12*). The hydrodynamic radius of the isolated 3'domain is not experimentally characterized, but we have calculated the radius of the 3'domain using following empirical formula published previously (*13*).

$$R_H = 5 \times 10^{-10} \times N^{0.38} \qquad -(2)$$

$R_H$ is the hydrodynamic radius, and N is the number of nucleotides.

The 475 nucleotides of the 3' major domain give a hydrodynamic radius of 52 Å. With a maximally possible separation of the Cy3-Cy5 FRET pair ($R_0$ = 51 Å; https://www.fpbase.org/fret/) on the hydrodynamic sphere of 104 Å (2x 52 Å), the FRET efficiency would drop to E= 0.014 based on the following formula:

$$E = 1/(1 + (R/R_0)^6) \qquad -(3)$$

E is FRET efficiency, $R_0$ is the Förster radius and R is the distance between donor and acceptor. In order to verify that all probe binding events are within FRET distance even in the unlikely events that the donor-acceptor pair is at the maximally possible separation, we repeated our experiments with the Cy3B-Cy5 dye combination ($R_0$ = 71.9 Å) which would give a FRET efficiency E=0.1 at 104 Å and thus, should be detected. Comparing our data for the Cy3-Cy5 and Cy3B-Cy5 FRET pairs shows an almost identical fraction of RNA molecule with probe binding for all three sites (fig. S2F). Thus, these experiments strongly support that we detect all probe binding events with our FRET-based detection approach.

Fraction of time the RNA remains bound by probe:

The heterogeneity observed in "fraction of time the RNA remains bound by probe" in the 3'domain can be explained by a heterogeneous population of RNA molecules, in which individual molecules transition between various RNA conformations. For example, the most heterogeneous H2829 region, in which "the fraction of time RNA remains bound by probe" varies by more than three orders of magnitude between different RNA molecules, is consistent with the idea that RNA molecules transition between H2829 accessible conformations (during which the probe can bind) and H2829 inaccessible conformations (in which S7 binds but not the probe). Thus, RNA molecules with little H2829 accessibility (e.g. having a single probe binding event) will have a much smaller "fraction of time the RNA remains bound by probe" than an RNA which is constantly accessible by the probe (which will have multiple probe binding events during the experimental time).

Increased H30 accessibility caused by secondary r-proteins:

In our previous work on the 3'domain assembly (*1*), and parallel work from the Woodson lab on the 5'domain assembly (*14*), we found that secondary r-proteins can chaperone nascent 3'domain rRNA folding by transiently interacting with the nascent RNA before the primary r-protein S7 is bound. At this stage, the binding sites for the secondary r-proteins are not created yet or only partially created, because secondary proteins can only stably bind to rRNA once the primary binding r-proteins are bound (*15, 16*). We believe that we cannot distinguish whether conformational selection or induced fit is responsible for the chaperoning effect mediated by the secondary r-proteins. The H30 accessible conformations belong to at least two types of RNA structures: 1) The S7+/H30+ class, which is increased in presence of secondary binding r-proteins, could represent a native assembly intermediate in which the H30 transiently opens (the H30 probe can bind) and closes (it needs to be closed when S7 binds). 2) The S7-/H30+ class, which does not increase in presence of secondary binding r-proteins, is likely non-native as it is kinetically trapped in an S7-binding incompetent conformation for at least 20-30 minutes (experimental time). From our data, we can tell that in the presence of secondary r-proteins, we see more molecules in the dynamic S7+/H30+ class (dynamic because it changes between H30 accessible and S7 binding competent conformations). However, whether this shift in population arises from conformational selection or induced fit mediated by the secondary r-proteins, a decision which is likely made as soon as the nascent binding sites emerge from the RNAP, cannot be distinguished from our data.

<u>SpikeTrain analysis is not applicable to our system:</u>

We originally considered using the SpikeTrain analysis, but were unsuccessful due to our much more complicated system. Walter and co-worker's SpikeTrain analysis was facilitated by their much simpler system with a smaller RNA molecule, which is present mainly in two conformations, and thus, allowed a simple interpretation of the data when the molecules switch from conformation 1 to conformation 2. In our case, the nascent RNA molecules are very heterogeneous and will switch between multiple different conformations. While our probe-only binding data (Fig. 2) may suggest that the RNA exists in two well-defined conformations, our more complex experiments, including, in addition, a protein (Fig. 3), show that this is not the case and that there are at least 4 conformational classes. Finally, even further extending our system by tracking accessibility of two sites with oligo binding and using a protein to assess the fold of the S7 binding site, we see that we have at least 8 conformational classes (Fig. 5). Therefore, using spike-train analysis on such data, which would report on the transition between two states, would be wrong data interpretation. Alternatively, the determination of the number of states from such heterogeneous data is extremely challenging and would require at least 1 order of magnitude more data (which is currently not feasible for such complex multi-color single-molecule experiments). Furthermore, the frequency of binding of our probe (to our much more structured rRNA as used in the study from Walter and co-workers) is too small to reliably track exact structural transitions. Thus, SpikeTrain analysis is not applicable to our system.

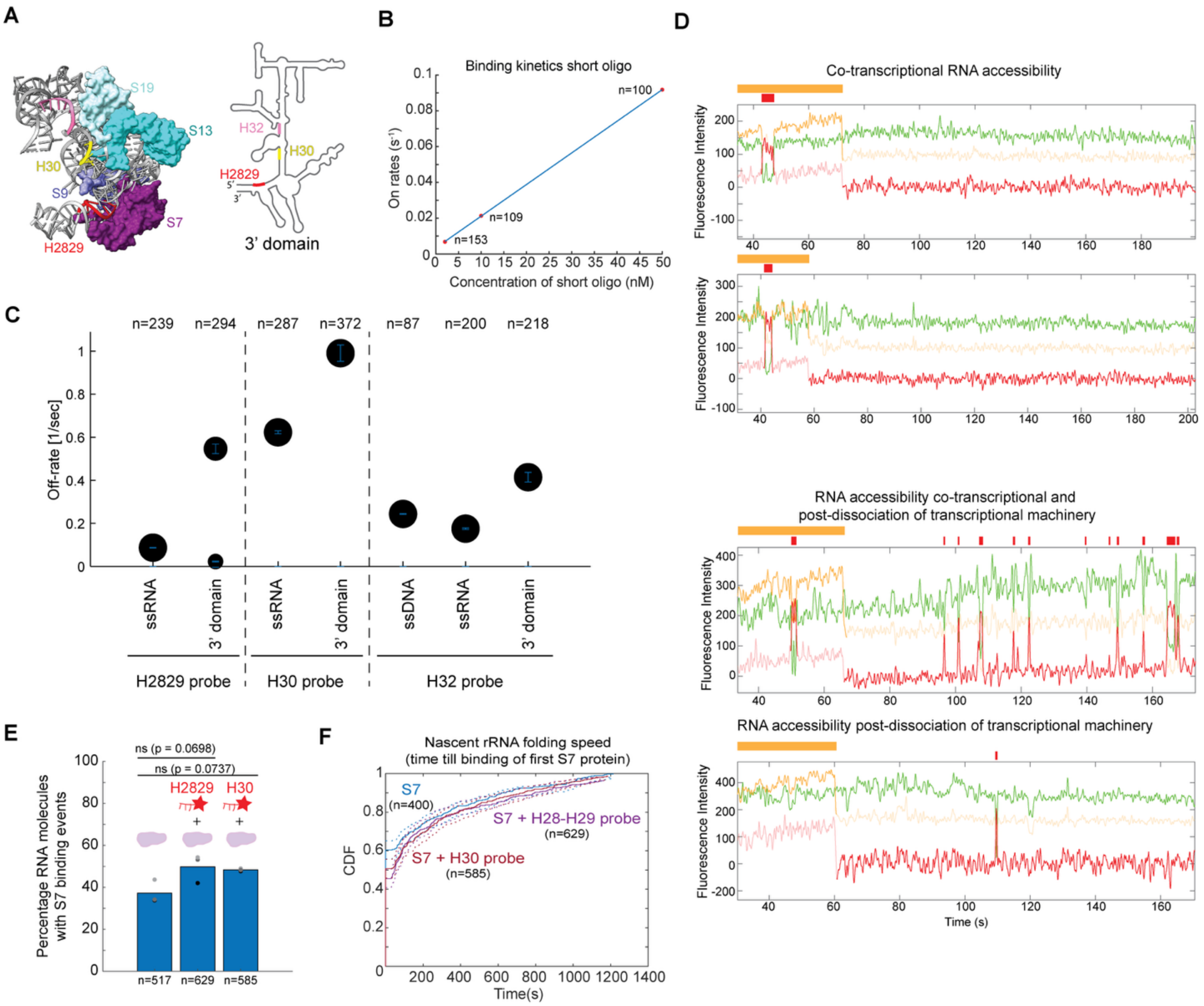


**Fig. S1: Characterization of probe binding**

**(A)** 3-dimensional structure (left) and secondary structure (right) of *E. coli* 30S ribosomal subunit zoomed into the S7 binding site. The regions probed in this study are shown: H2829 (red), H30 (yellow), H32 (pink) and surfaces of r-proteins S7 (purple), S9 (blue), S19 (sea blue) and S13 (cyan) (PDB accession code: 4V9P). **(B)** On-rate of the short DNA probe binding to H2829 ssRNA. **(C)** Comparison of the off-rate of the probes to the ssRNA, 3'domain and ssDNA. **(D)** Example smoothed traces showing binding of the H30 probe co-transcriptionally and post dissociation of the transcription machinery. Simplified representation of H30 probe binding events (red), and transcription signal (yellow) are shown above the trace. **(E)** The percentage of RNA molecules binding S7 (purple) in presence of the different short DNA probes (bars represent means of replicates). P-values (p) from two-sample Student's t-test (unequal variances): ns represents non-significant. **(F)** The cumulative distribution of the time from transcription start till appearance of the first S7 binding event in absence and presence of different short DNA probes (each curve is plotted by pooling data from 2 or 3 replicates, the dotted lines represent confidence bounds of the CDF fit). **(B,C,E,F)** Number of molecules analyzed (n) are shown.

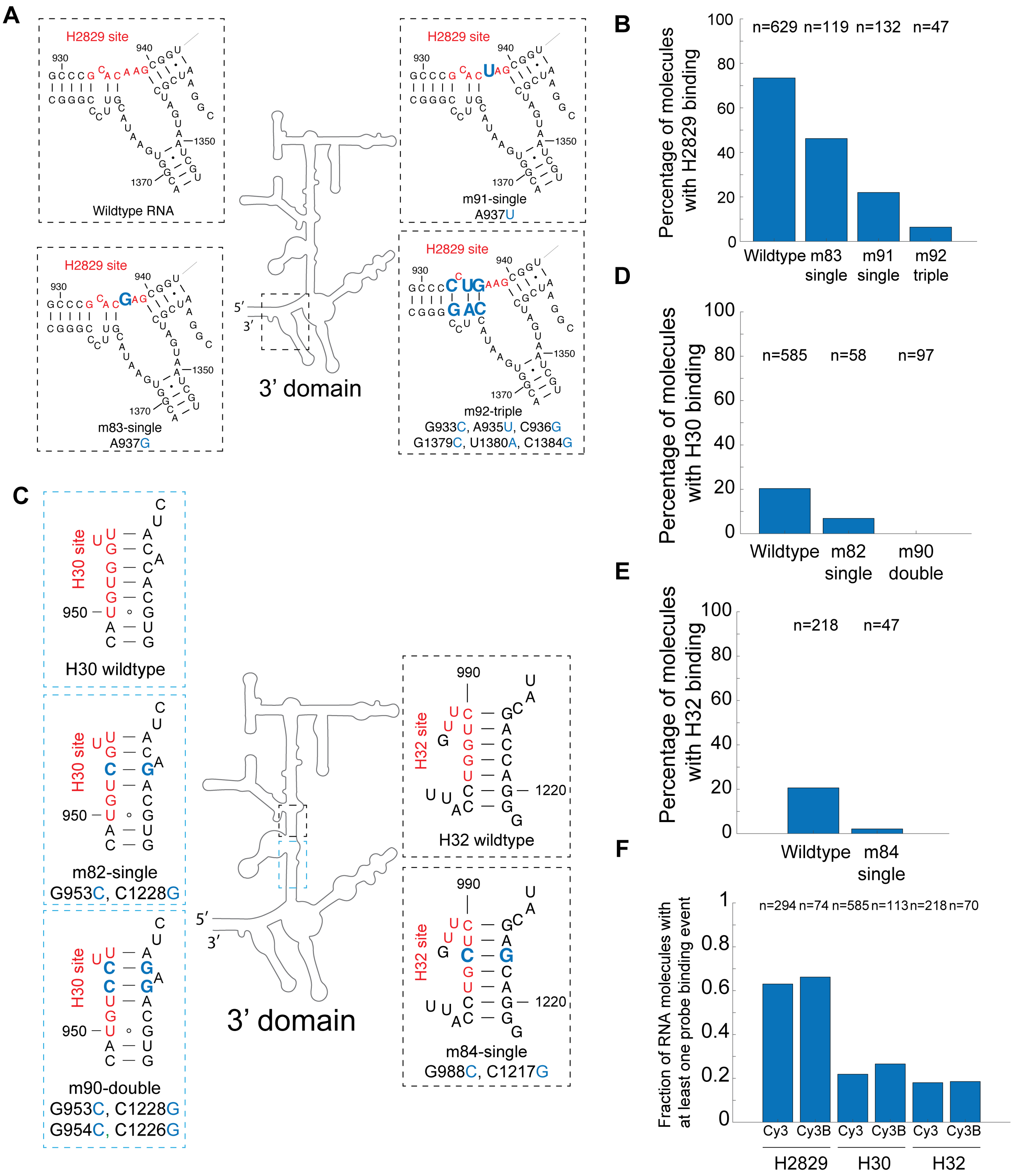


**Fig. S2: Probe binding is abolished when target sites on the 3’domain are mutated.** Schematic of 3’domain indicating the single, double or triple compensatory point mutations introduced (blue) and the target sites (red) for **(A)** H2829, **(C)** H30 and H32 target sites. Percentage of molecules accessible at **(B)** H2829, **(D)** H30, and **(E)** H32 sites to different constructs. **(F)** Comparison of the fraction of molecules with probe binding events to the 3’domain with Cy3 or Cy3B dye as donor and Cy5 dye as acceptor. Number of molecules analyzed (n) are shown. The data for wildtype (in **B,D,E**) and Cy3 (in **F**) is same as presented in Fig. 2B.

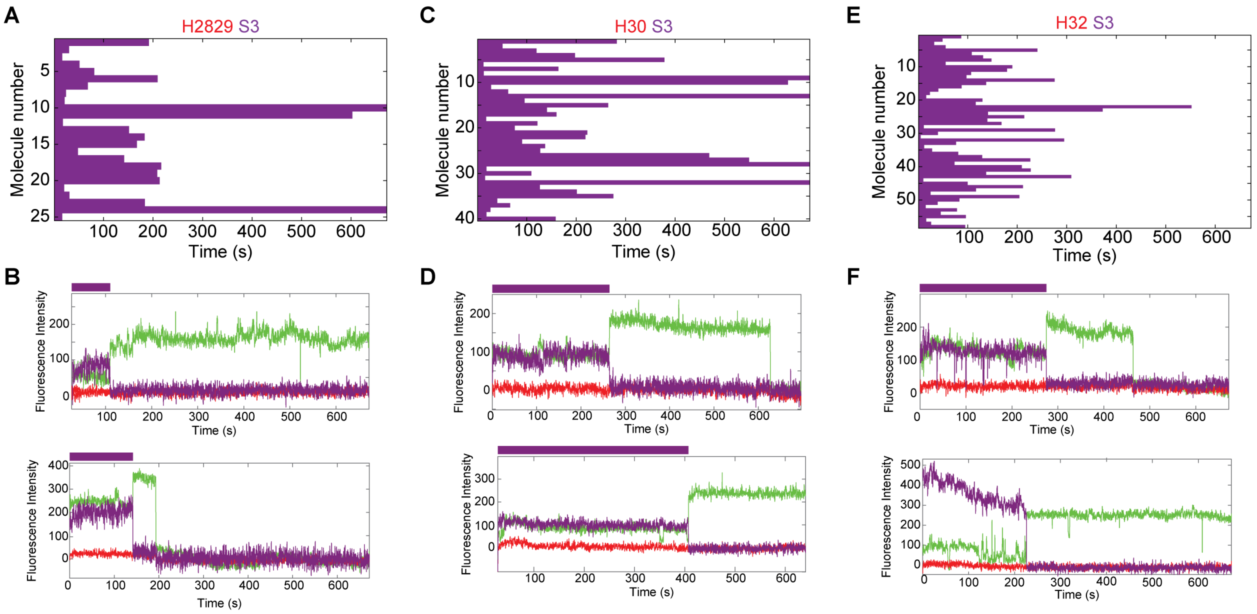


**Fig. S3: RNA is inaccessible to probes in fully assembled 3'domain complex.**
Probing RNA accessibility of the assembled 3'domain complex at **(A)** H2829, **(C)** H30 and **(E)** H32 sites, respectively. Example smoothed traces show bound S3 (purple – Cy5.5 dye), which is a marker for the fully assembled 3'domain (*1*). We did not detect probe binding (expected in red – Cy5 dye) to the subset of molecules with bound S3 for **(B)** H2829, **(D)** H30 and **(F)** H32 sites, respectively.

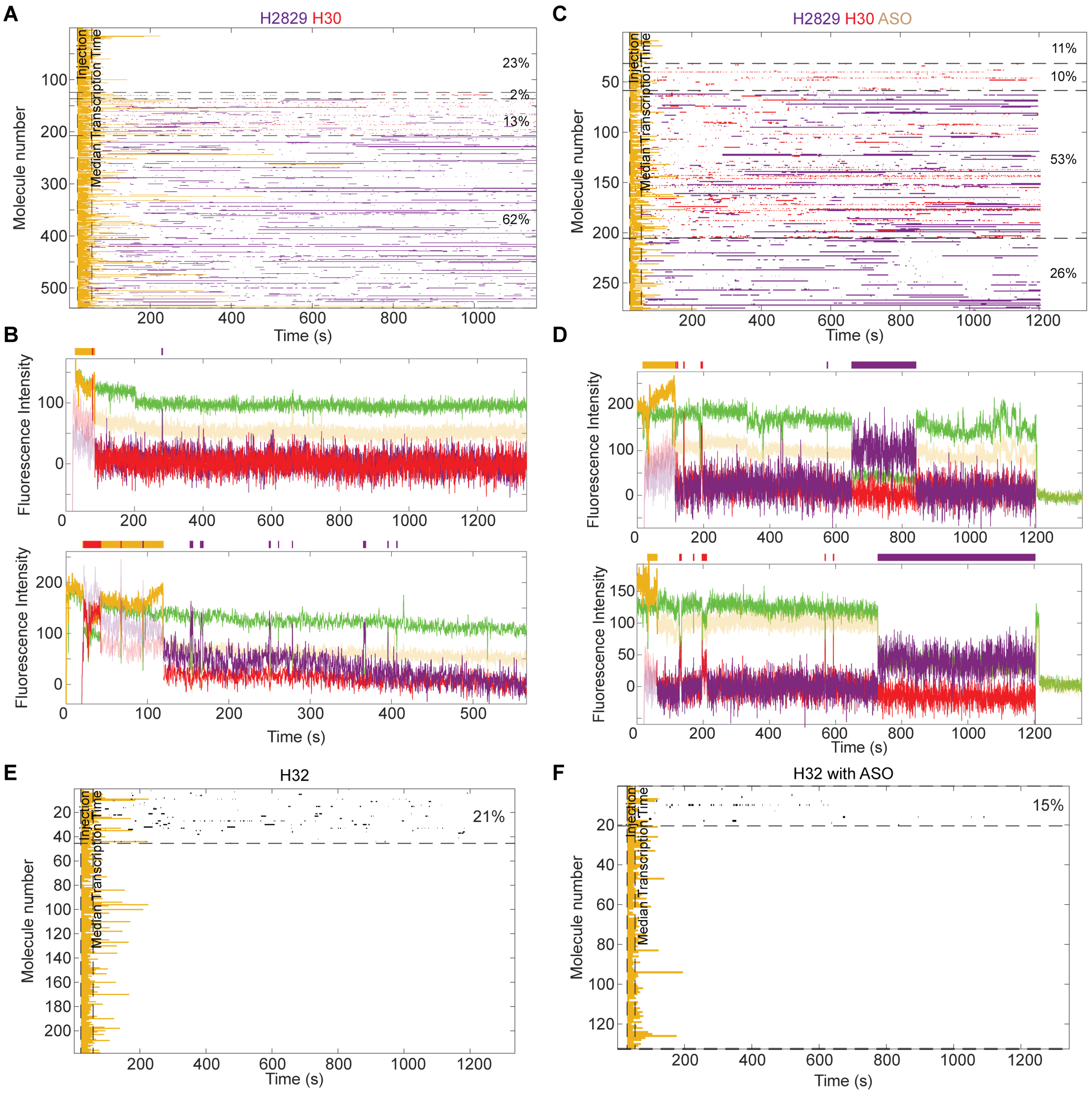


**Fig. S4: Effect of ASOs on the RNA accessibility for different regions of the 3'domain rRNA. (A,C)** Rasterplot showing individual molecules as rows and **(B,D)** smoothed example traces with transcription signal (yellow – Cy3.5 dye), H30 DNA probe (red – Cy5 dye) and H2829 DNA probe (purple – Cy5.5 dye) binding events shown as colored bars in absence **(A)** and presence of ASOs **(C)**, respectively**. (E)** Rasterplots showing individual molecules as rows with black representing H32 probe binding in absence **(E)** and presence of ASOs **(F)**. **(A,C,E)** plotted by pooling 2 replicates, **(F)** one experiment**.**

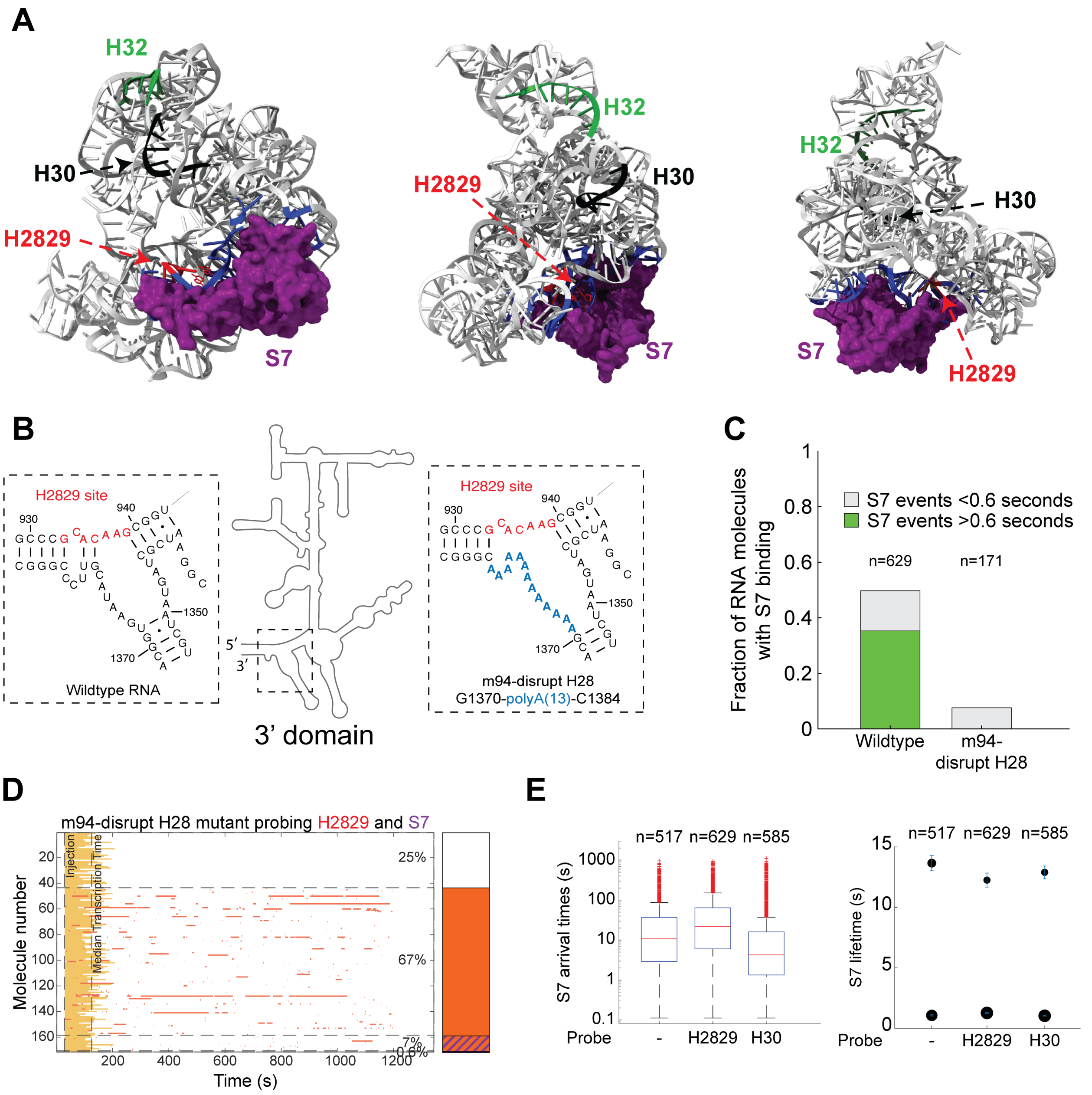


**Fig. S5: RNA mutations strongly reduce S7 binding**

**(A)** 3-dimensional structure of 3’domain bound by S7 from different angles. The regions probed in this study are shown: H2829 (red), H30 (black), H32 (green) and surfaces of r-proteins S7 (purple), contact sites of S7 to the RNA (blue) (PDB accession code: 4V9P). **(B)** Schematic of mutations on the RNA showing probe binding site (red) and mutations (blue). **(C)** Fraction of RNA molecules with S7 binding to wildtype and mutant RNA. Wildtype data is the same as presented in fig. S1E. **(D)** Rasterplot of m94-H28 disrupting mutant showing transcription (yellow – Cy3.5 dye), H2829 probe binding (red – Cy5 dye) and almost no S7 binding (purple – Cy5.5 dye). **(E)** S7 arrival times (left) and lifetimes (right) at 20 nM concentration of S7-Cy5.5 to the wildtype 3’domain RNA. Number of molecules analyzed (n) are shown.

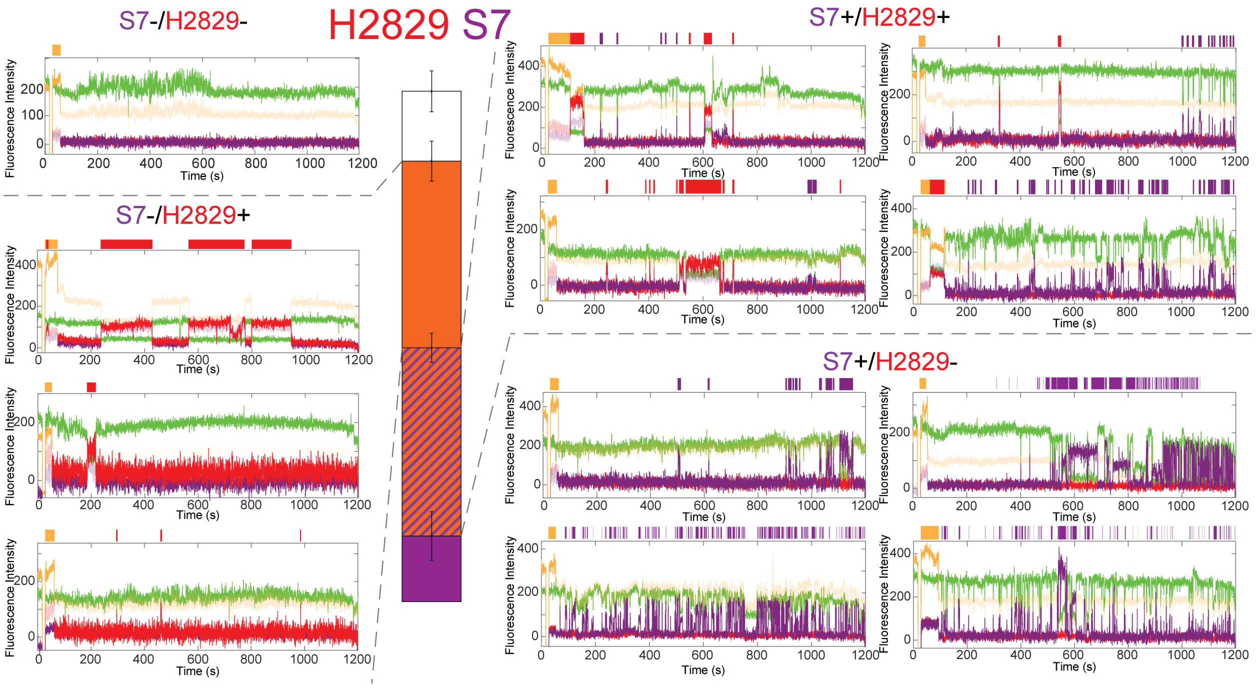


**Fig. S6: H2829 accessibility and S7 binding is highly heterogeneous.**
Example smoothed traces across different nascent RNA folding classes showing transcription (yellow – Cy3.5 dye), H2829 accessibility (red – Cy5 dye) and S7 binding (purple – Cy5.5 dye).

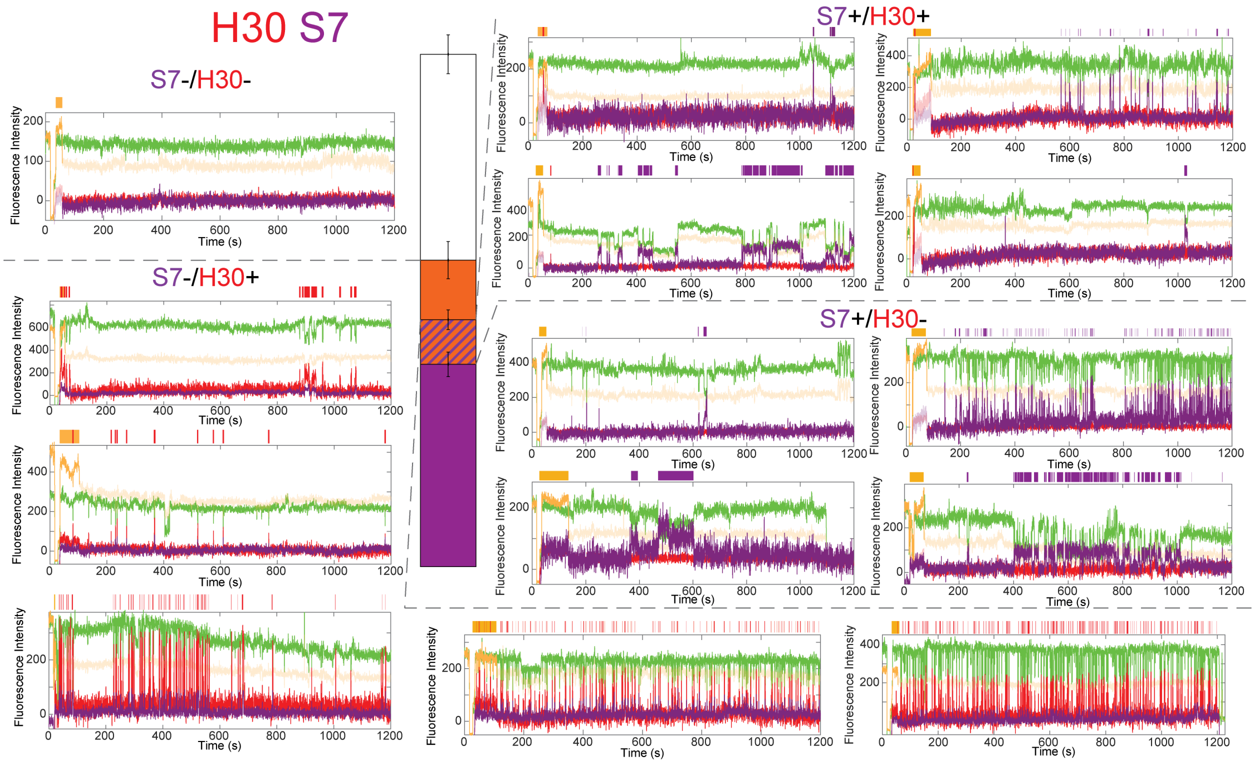


**Fig. S7: H30 accessibility and S7 binding is highly heterogeneous.**
Example smoothed traces across different nascent RNA folding classes showing transcription (yellow – Cy3.5 dye), H30 accessibility (red – Cy5 dye) and S7 binding (purple – Cy5.5 dye).

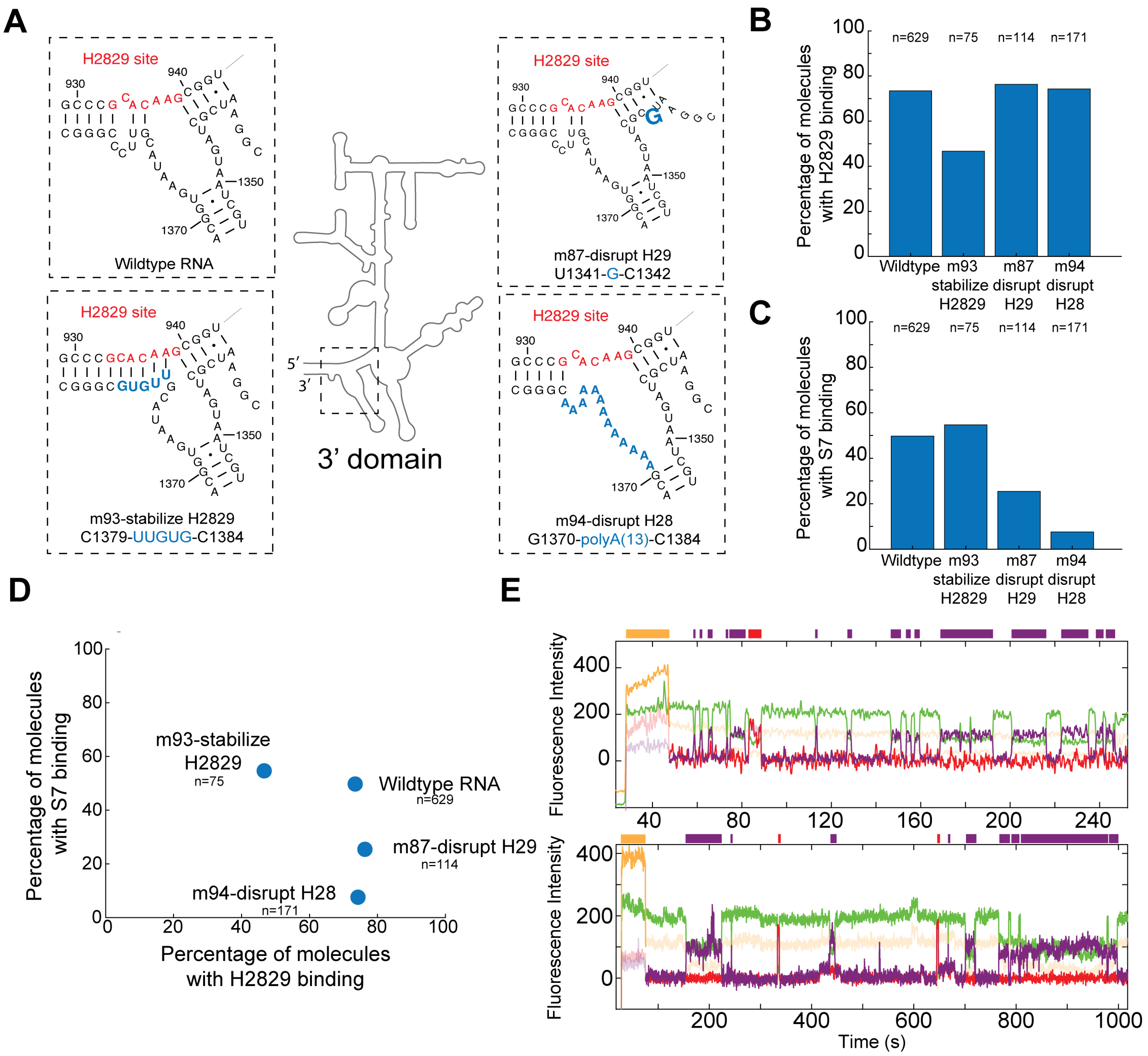


**Fig. S8: Mutations on the H2829 region of the 3’domain shift the dynamic equilibrium.** **(A)** Schematic of mutations on the RNA showing probe binding site (red) and mutations (blue). Percentage of molecules with **(B)** H2829 accessibility and **(C)** S7 binding in different constructs. Wildtype and m94 disrupting H28 mutant data in **(B,C,D)** is the same as presented in Fig. 3G and fig. S5, respectively. **(D)** A scatter plot represents the relative changes observed in percentage molecules accessible at H2829 accessibility and S7 bound across different constructs. Number of molecules analyzed (n) in **(B-D)** are shown. **(E)** Example smoothed traces showing transcription (yellow – Cy3.5 dye) and dynamic equilibrium between H2829 being accessible (red – Cy5 dye) and S7 bound (purple – Cy5.5 dye).

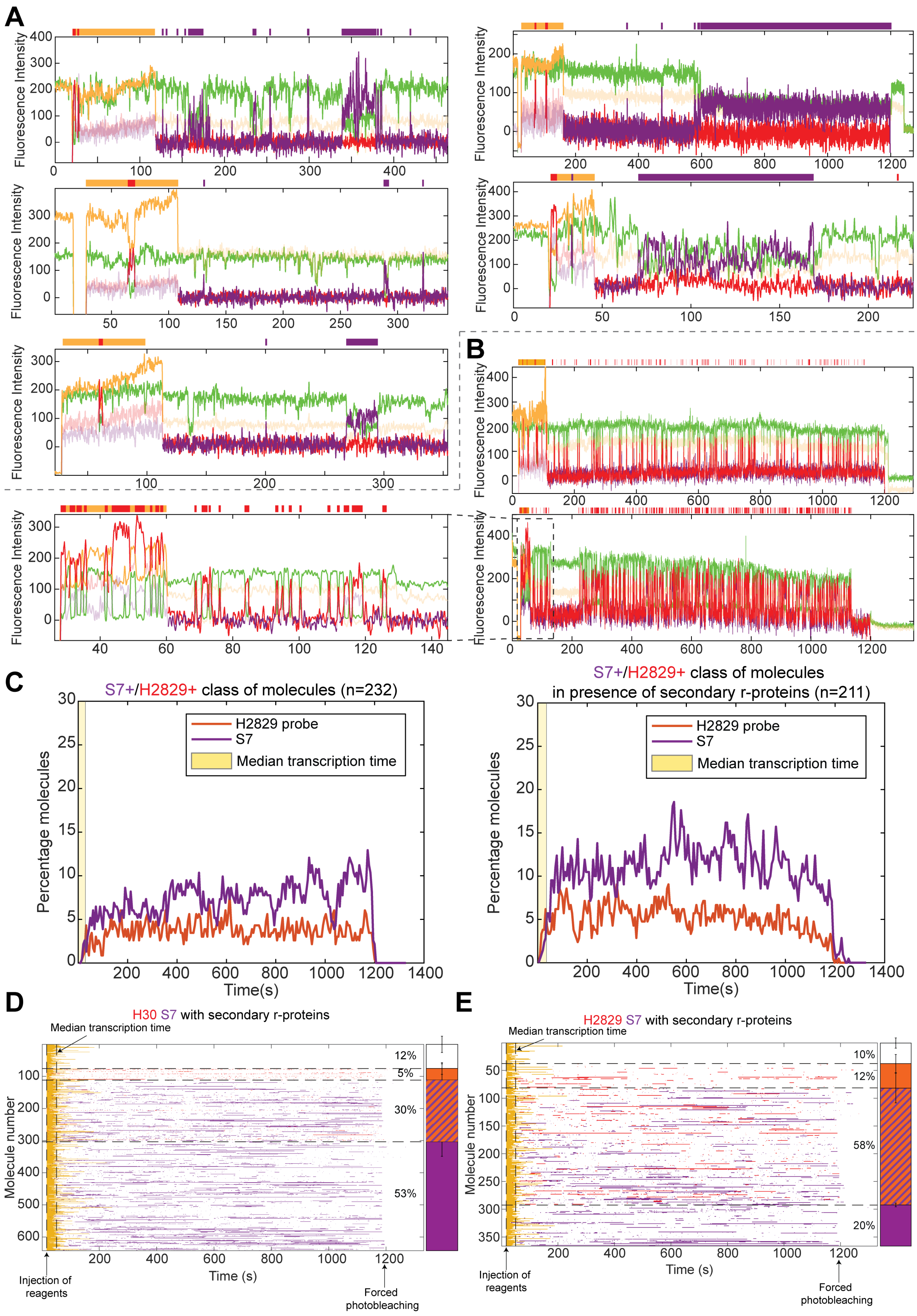
A
B
C
D
E
Fluorescence Intensity
S7+/H2829+ class of molecules (n=232)
S7+/H2829+ class of molecules
in presence of secondary r-proteins (n=211)
H2829 probe
S7
Median transcription time
Percentage molecules
Time(s)
H30 S7 with secondary r-proteins
H2829 S7 with secondary r-proteins
Median transcription time
Molecule number
Time (s)
Injection of reagents
Forced photobleaching
12%
5%
30%
53%
10%
12%
58%
20%

**Fig. S9: Secondary r-proteins have variable effect on the different classes and on different regions of the RNA.** Example smoothened traces of class **(A)** S7+/H30+ and **(B)** S7-/H30+ showing transcription (yellow – Cy3.5 dye), H30 probe binding (red – Cy5 dye) and S7 binding (purple – Cy5.5 dye) in the presence of secondary r-proteins. **(C)** Time dependent analysis of the S7+/H2829+ class of molecules in absence (left) and presence of the secondary r-proteins (right). The region highlighted in yellow represents the median time RNA is associated to transcription machinery. Number of molecules analyzed (n) are shown. **(D,E)** Rasterplots showing the individual molecules as rows with transcription (yellow), DNA probe (red) and S7 (purple) binding events in presence of secondary r-proteins, shown as colored bars for H30 **(D)** and H2829 **(E)** sites, respectively. All data plotted by pooling 3 replicates. Error bars show weighted standard deviations.

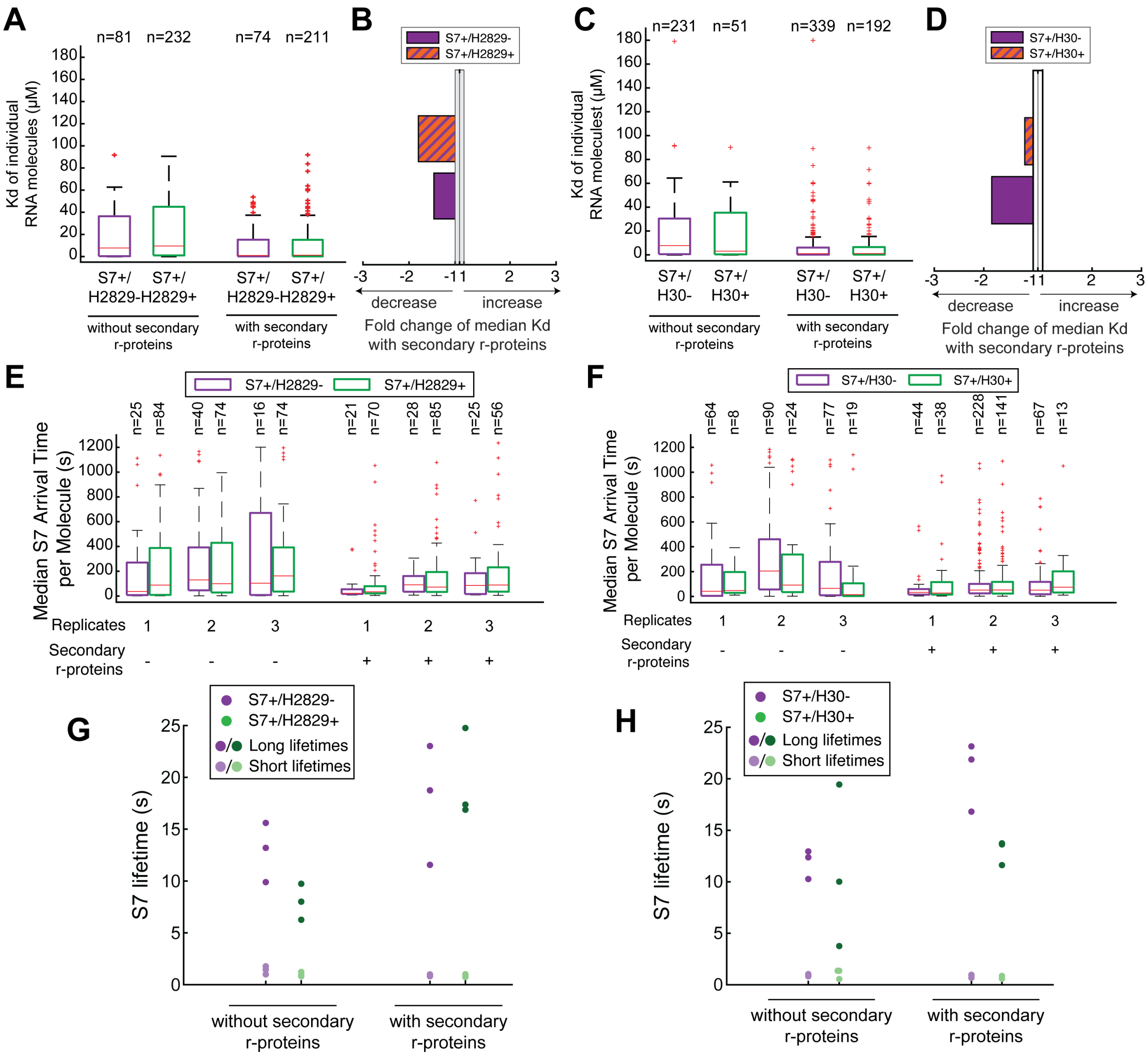


**Fig. S10: Thermodynamics and kinetics of S7 across RNA folding classes.**
**(A,C)** Boxplot of $K_d$ for S7 of individual molecules calculated between 200 s and photobleaching and **(B,D)** fold change across nascent RNA folding classes in the absence and presence of secondary r-proteins for **(A,B)** H2829 site and **(C,D)** H30 site. **(E,F)** Median arrival time per molecule and **(G,H)** fits of S7-bound dwells to two exponential function across nascent RNA folding classes in the absence and presence of secondary r-proteins for **(E,G)** H2829 site and **(F,H)** H30 site. Different dots of same color represent replicates and shades of color represent different populations of S7 lifetimes in **(G,H)**. Number of molecules analyzed (n) are shown.

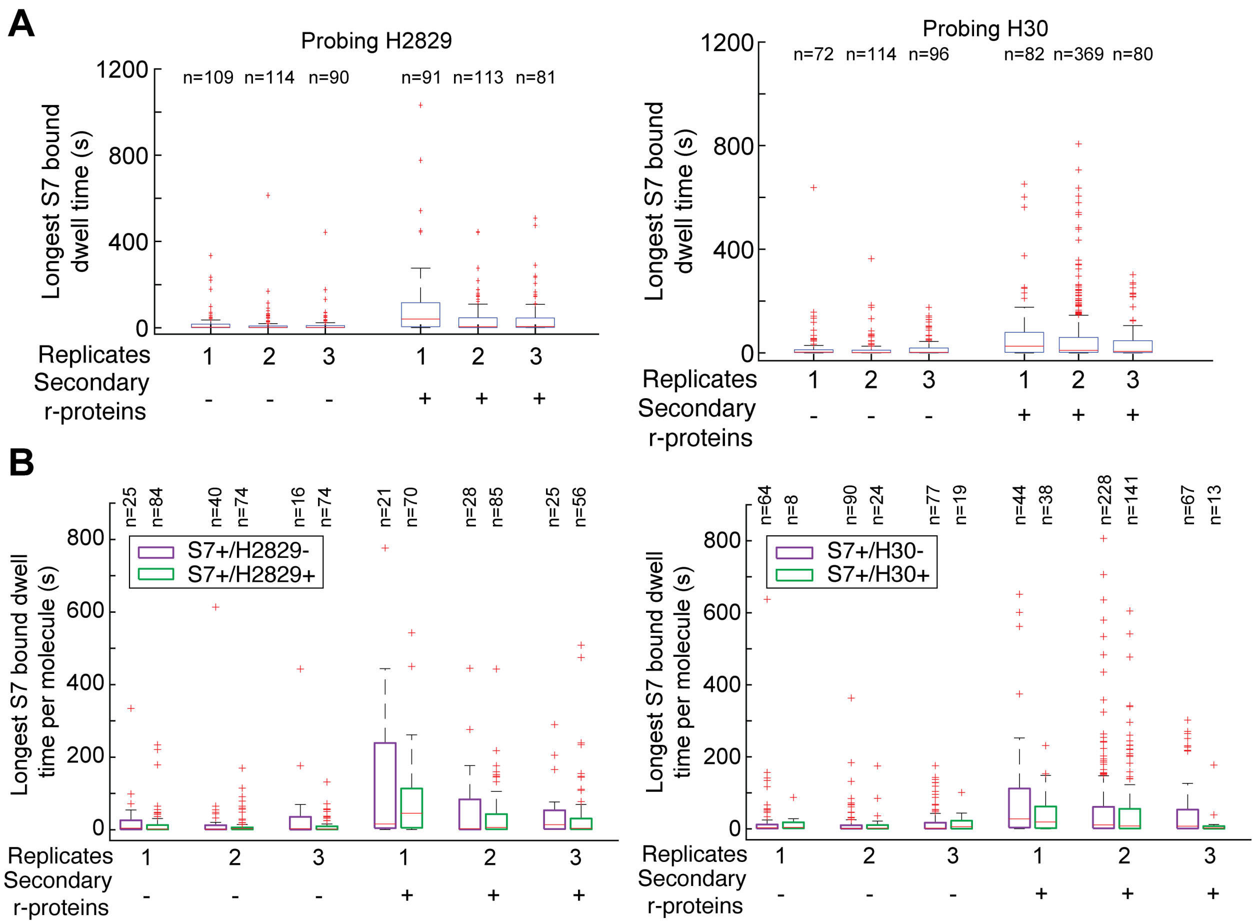


**Fig. S11: Secondary r-proteins increase the longest S7 bound lifetime across all classes.** Boxplot of **(A)** the longest S7 binding events of all the molecules, and **(B)** of molecules in individual classes for experiment probing H2829 (left) and H30 (right). Number of molecules analyzed (n) are shown. Plotting the longest S7 event per trace, instead of fitting all S7-bound dwell times (like in fig. S10), allows enrichment of events in which the complexes are assembled with secondary r-proteins. See more details in (*1*).

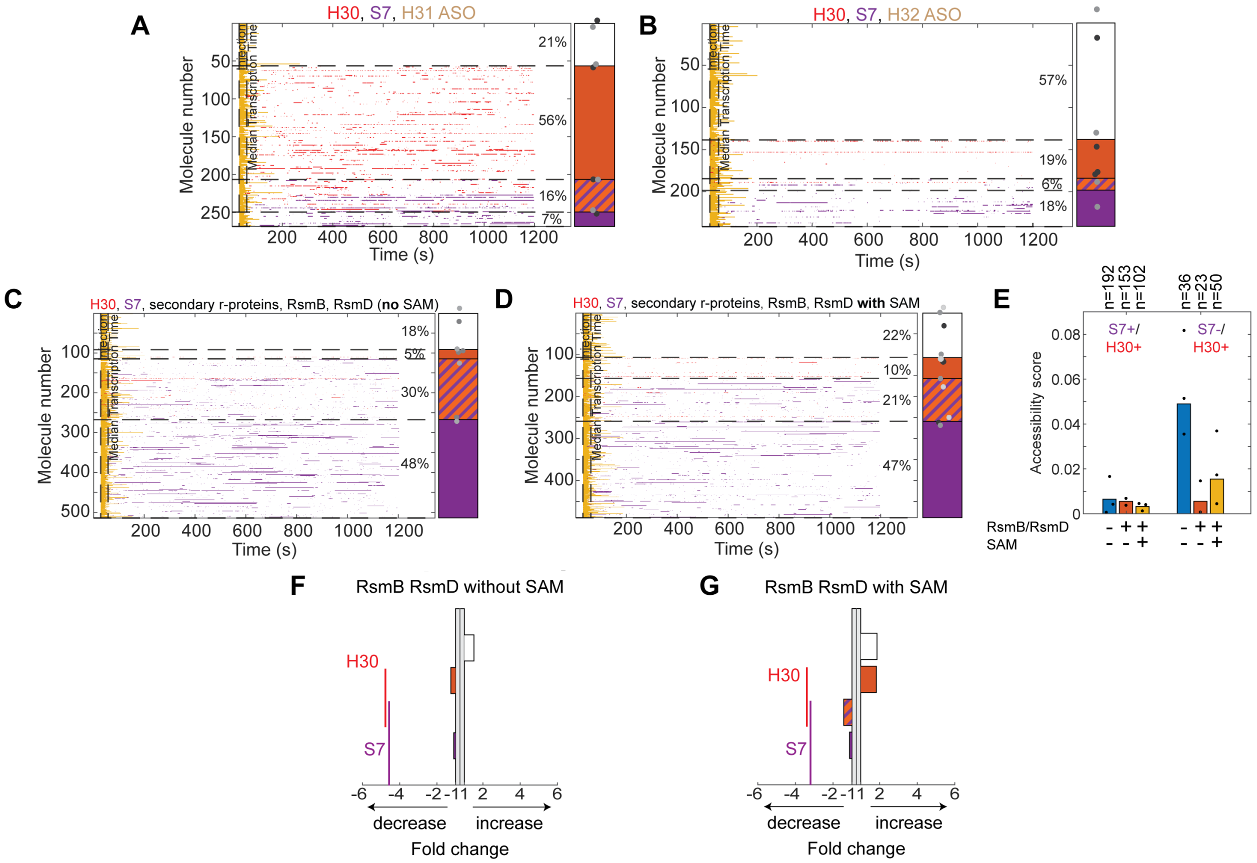


**Fig. S12: Quantifying the modulation effect by ASOs and rRNA modification enzymes**

**(A-D)** Rasterplots showing individual molecules as rows for probing H30 region (red) and S7 binding (purple) on the 3'domain in presence of the H31 ASO **(A)** or H32 ASO **(B)** (both in absence of secondary r-proteins); or RsmB, RsmD without SAM **(C)** or with SAM **(D)** (both in presence of secondary r-proteins). Stacked bars are weighted means of replicates shown as shades of grey dots. **(E)** H30 accessibility score in RNA folding classes in presence of only secondary r-proteins (blue), in addition with RsmB and RsmD (orange) and in addition in presence of SAM (yellow). Mean accessibility score calculated by pooling 2 or 3 replicates. Dots represent individual replicates, number of molecules analyzed (n) are shown. **(F,G)** Relative fold-change in RNA class distributions upon addition of RsmB/RsmD in absence **(F)** or presence **(G)** of SAM (always in presence of S9, S13 and S19); data from Fig. 4E.

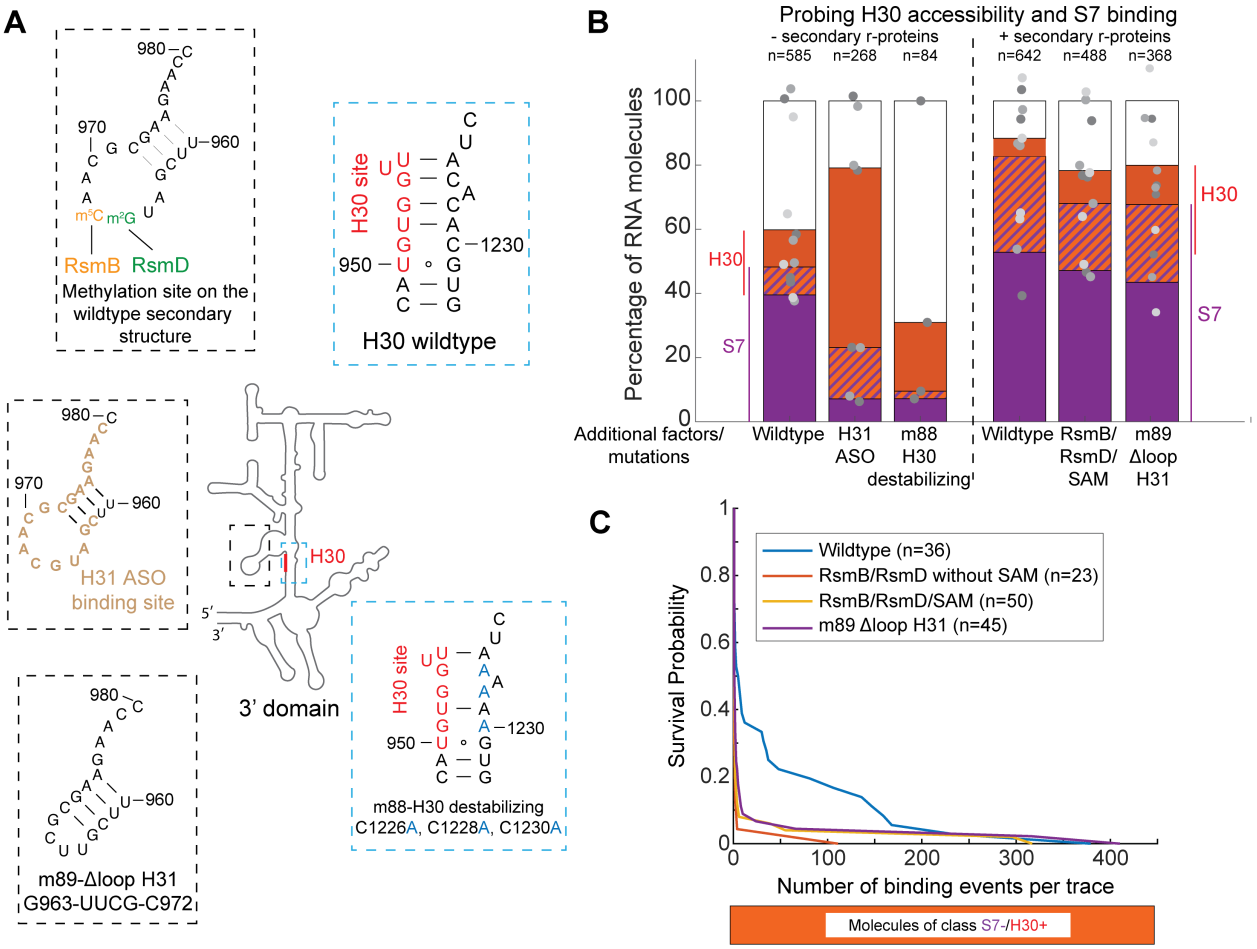


**Fig. S13: H30 region is sensitive to mutations and RNA modification enzyme binding**
**(A)** Schematic of mutations and ligand binding sites on the RNA. **(B)** Percentage of molecules in RNA folding classes affected by various perturbations. **(C)** Comparison of survival probability of the number of H30 binding events per trace in the S7-/H30+ class in the presence of secondary r-proteins. Dots represent replicates in **(B)** and n represents number of molecules. Data for wildtype with and without secondary r-proteins, and H31 ASO, RsmB/RsmD without and with SAM conditions is the same as presented in Fig. 3G and Fig. 4B,E, respectively.

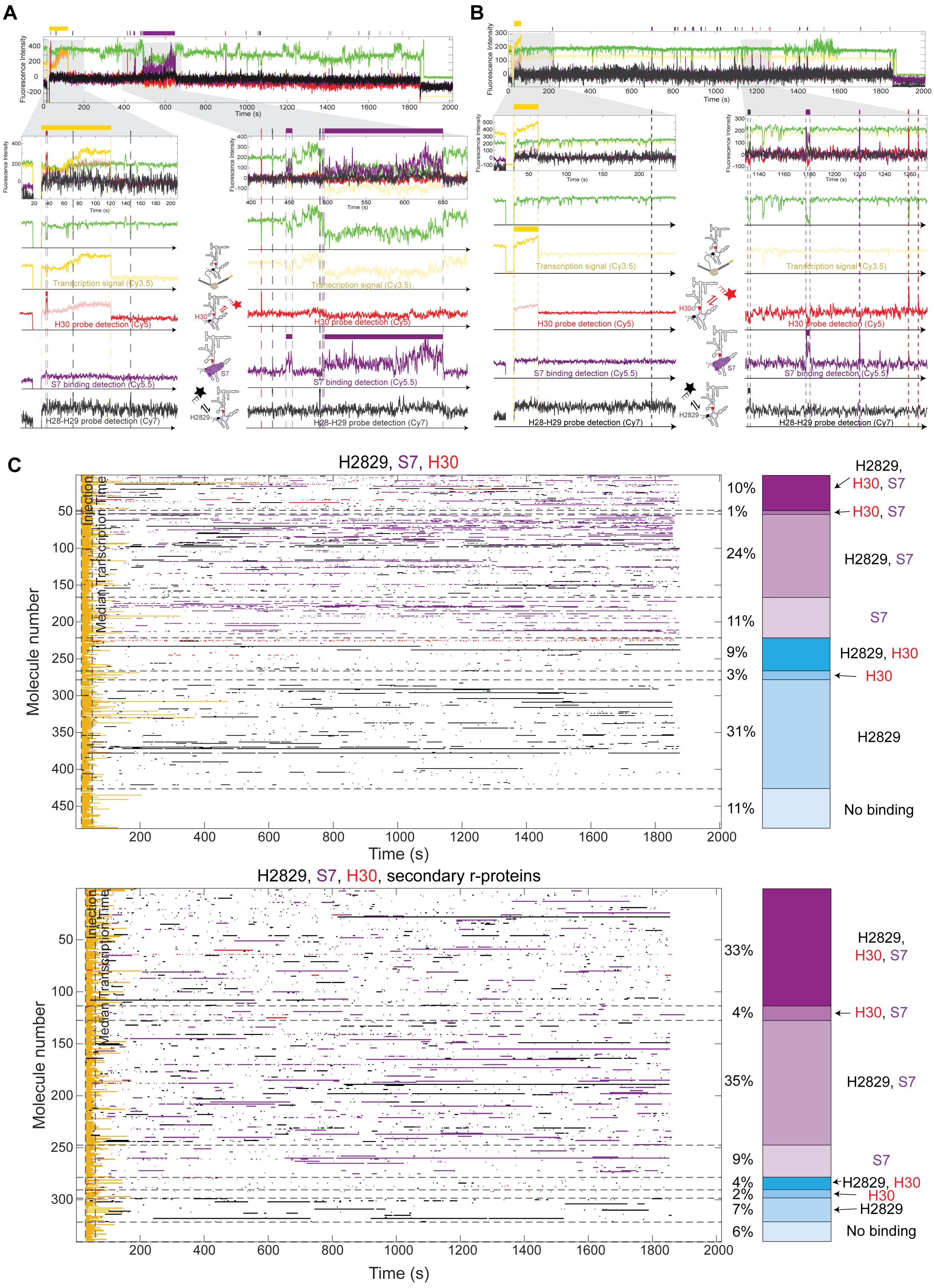
A
B
C
Fluorescence Intensity
Time (s)
Transcription signal (Cy3.5)
H30 probe detection (Cy5)
S7 binding detection (Cy5.5)
H28-H29 probe detection (Cy7)
H30
S7
H2829
H2829, S7, H30
Molecule number
Injection
Median Transcription Time
10%
1%
24%
11%
9%
3%
31%
11%
H2829, H30, S7
H30, S7
H2829, S7
S7
H2829, H30
H30
H2829
No binding
H2829, S7, H30, secondary r-proteins
33%
4%
35%
9%
4%
2%
7%
6%

**Fig. S14: Multi-site probing experiments show heterogeneity in RNA folding**

**(A,B)** Example smoothed traces of the 5-color data showing FRET donor (green – Cy3 dye), transcription signal (yellow – Cy3.5 dye), H30 probe (red – Cy5 dye), S7 r-protein (purple – Cy5.5 dye) and H2829 probe binding (black – Cy7 dye). Simplified representation of binding events is shown on top of the traces as bars. **(C)** Rasterplots of the multi-site probing experiments where individual molecules are rows showing transcription (yellow), binding of DNA probes to H30 (red) and H2829 (black), and r-protein S7 (purple) in absence (left) and presence of secondary r-proteins (right). Plotted by pooling 4 replicates – without, 2 replicates – with secondary r-proteins.

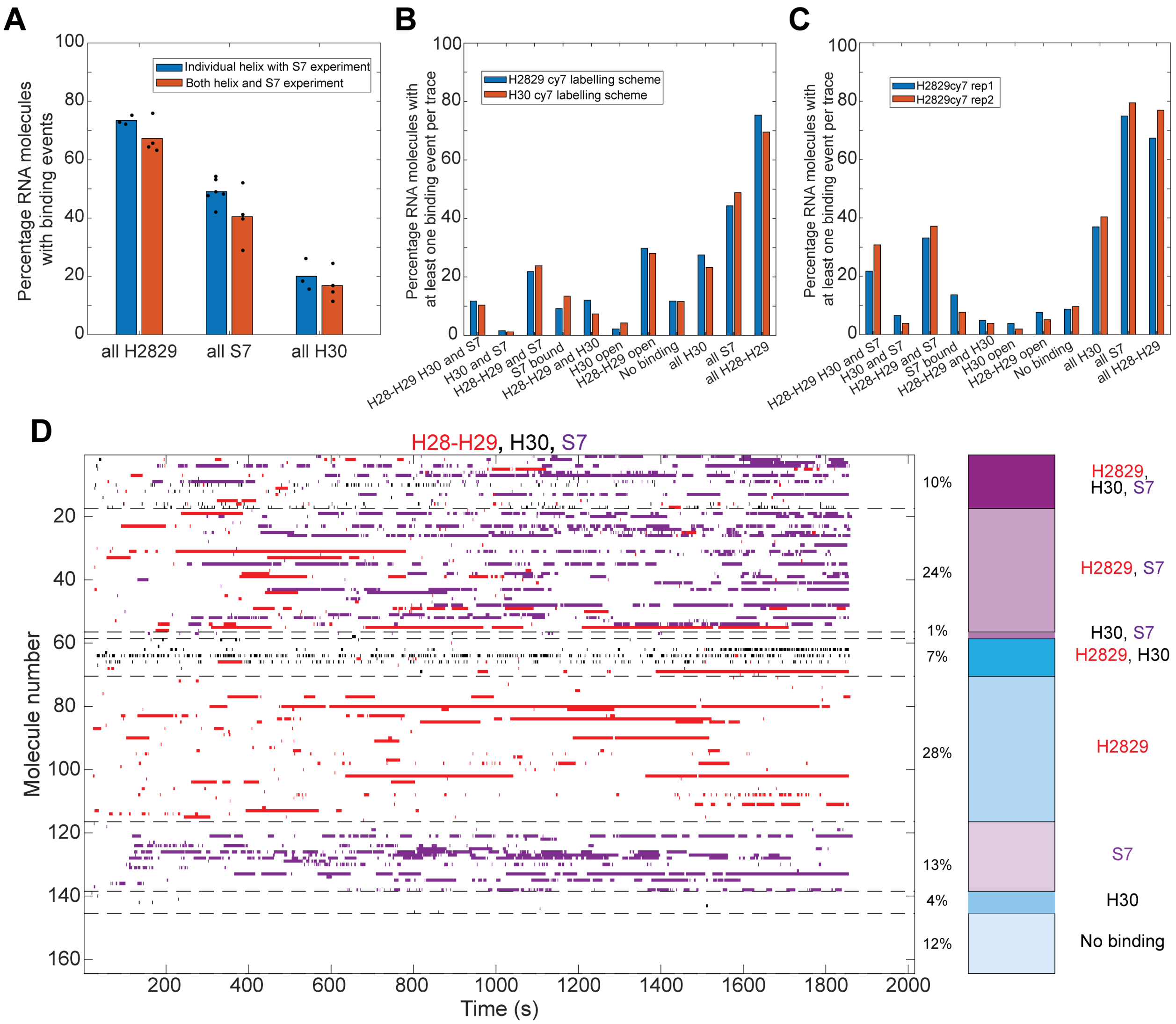


**Fig. S15: Comparing multi-site probing experiments with single-site probing experiments**
**(A)** Percentage of molecules that show probe or S7 binding events in individual or two-site probing experiment in absence of secondary r-proteins. Bars represent mean of replicates shown as dots. **(B)** Comparing replicates using different dye labelling schemes in absence of secondary r-proteins (blue bars: Cy7-H2829-DNA probe, Cy5-H30-DNA probe and Cy5.5-S7; red bars: Cy5-H2829-DNA probe, Cy7-H30-DNA probe and Cy5.5-S7). **(C)** Comparing replicates for two-site probing with S7 binding in presence of secondary r-proteins (replicates are colored in blue and red). **(D)** Rasterplot of multi-site experiment where individual molecules are rows showing binding of DNA probe to H2829 (red), H30 (black) and S7 binding (purple) to the 3'domain in absence of secondary r-proteins; H2829-Cy5 dye, H30-Cy7 dye, S7-Cy5.5 dye labeling scheme.

| Dataset | Koff,1 | Koff,2 | Percentage events described by koff,1 |
|---|---|---|---|
| Duss et. al, Cell, 2019 | 0.79 ± 0.03 s-1 | 0.046 ± 0.004 s-1 | 75% |
| S7 | 0.94 ± 0.0045 s-1 | 0.073 ± 0.0035 s-1 | 63% |
| S7 + H2829 probe | 0.79 ± 0.025 s-1 | 0.082 ± 0.0035 s-1 | 76% |
| S7 + H30 probe | 0.97 ± 0.025 s-1 | 0.077 ± 0.0035 s-1 | 76% |

**Table S1: Comparison of S7 off-rates from this study to previous study.**

**Materials**

Overview of artificial DNA sequences used.

Backbone of the DNA template used.
RNAP promoter (green), hybridization sequence for immobilization of stalled complex through 5'-end of nascent RNA (red), binding site for DNA oligo binding to 5'end of nascent RNA (blue), 3'domain sequence (see further below), binding site for labeled DNA oligo binding to 3'-end of nascent RNA (orange) and triple transcription terminator (black).

The DNA sequence is:
ctggcagttt taggctgatt tggttgaatg ttgcgcggtc agaaaattat tttaaatttc ctcttgtcag gccggaataa ctccctataa
tgcgccacc ACT ACCAC CACCC AACCA ACACA CC AAC CAC TCC AAT TAC ATA CAC
C (3'domain) CCCTA TCCCT TATCT TAAC
GGCTCCTTTTGGAGCCTTTTTTTTTGGAGATTTTCTAAAACGAAAGGCTCAGT
CGAAAGACTGGGCCTTTCGTTTTATCT TAATC ACACT GGCTC ACCTT CGGGT
GGGCC TTTCT GCGTT TAT

3’domain (5' to 3'):
tgacgggggc ccgcacaagc ggtggagcat gtggtttaat tcgatgcaac gcgaagaacc ttacctggtc ttgacatcca
cggaagtttt cagagatgag aatgtgcctt cgggaaccgt gagacaggtg ctgcatggct gtcgtcagct cgtgttgtga
aatgttgggt taagtcccgc aacgagcgca acccttatcc tttgttgcca gcggtccggc cgggaactca aaggagactg
ccagtgataa actggaggaa ggtggggatg acgtcaagtc atcatggccc ttacgaccag ggctacacac gtgctacaat
ggcgcataca aagagaagcg acctcgcgag agcaagcgga cctcataaag tgcgtcgtag tccggattgg agtctgcaac
tcgactccat gaagtcggaa tcgctagtaa tcgtggatca gaatgccacg gtgaatacgt tcccgggcct tgtaca

m82 (5' to 3'):
tgacgggggcccgcacaagcggtggagcatgtCgtttaattcgatgcaacgcgaagaaccttacctggtcttgacatccacggaagttttc
agagatgagaatgtgccttcgggaaccgtgagacaggtgctgcatggctgtcgtcagctcgtgttgtgaaatgttgggttaagtcccgcaac
gagcgcaacccttatcctttgttgccagcggtccggccgggaactcaaaggagactgccagtgataaactggaggaaggtggggatgac
gtcaagtcatcatggcccttacgaccagggctacaGacgtgctacaatggcgcatacaaagagaagcgacctcgcgagagcaagcgga
cctcataaagtgcgtcgtagtccggattggagtctgcaactcgactccatgaagtcggaatcgctagtaatcgtggatcagaatgccacggt
gaatacgttcccgggccttgtaca

m83 (5' to 3'):
tgacgggggcccgcacGagcggtggagcatgtggtttaattcgatgcaacgcgaagaaccttacctggtcttgacatccacggaagttttc
agagatgagaatgtgccttcgggaaccgtgagacaggtgctgcatggctgtcgtcagctcgtgttgtgaaatgttgggttaagtcccgcaac
gagcgcaacccttatcctttgttgccagcggtccggccgggaactcaaaggagactgccagtgataaactggaggaaggtggggatgac
gtcaagtcatcatggcccttacgaccagggctacacacgtgctacaatggcgcatacaaagagaagcgacctcgcgagagcaagcggac
ctcataaagtgcgtcgtagtccggattggagtctgcaactcgactccatgaagtcggaatcgctagtaatcgtggatcagaatgccacggtg
aatacgttcccgggccttgtaca

m84 (5' to 3'):
tgacgggggcccgcacaagcggtggagcatgtggtttaattcgatgcaacgcgaagaaccttacctgCtcttgacatccacggaagttttc
agagatgagaatgtgccttcgggaaccgtgagacaggtgctgcatggctgtcgtcagctcgtgttgtgaaatgttgggttaagtcccgcaac
gagcgcaacccttatcctttgttgccagcggtccggccgggaactcaaaggagactgccagtgataaactggaggaaggtggggatgac

gtcaagtcatcatggcccttacgaGcagggctacacacgtgctacaatggcgcatacaaagagaagcgacctcgcgagagcaagcggacctcataaagtgcgtcgtagtccggattggagtctgcaactcgactccatgaagtcggaatcgctagtaatcgtggatcagaatgccacggtgaatacgttcccgggccttgtaca

m87 (5' to 3'):
tgacgggggcccgcacaagcggtggagcatgtggtttaattcgatgcaacgcgaagaaccttacctggtcttgacatccacggaagttttcagagatgagaatgtgccttcgggaaccgtgagacaggtgctgcatggctgtcgtcagctcgtgttgtgaaatgttgggttaagtcccgcaacgagcgcaacccttatcctttgttgccagcggtccggccgggaactcaaaggagactgccagtgataaactggaggaaggtggggatgacgtcaagtcatcatggcccttacgaccagggctacacacgtgctacaatggcgcatacaaagagaagcgacctcgcgagagcaagcggacctcataaagtgcgtcgtagtccggattggagtctgcaactcgactccatgaagtcggaatGcgctagtaatcgtggatcagaatgccacggtgaatacgttcccgggccttgtaca

m88 (5' to 3'):
tgacgggggcccgcacaagcggtggagcatgtggtttaattcgatgcaacgcgaagaaccttacctggtcttgacatccacggaagttttcagagatgagaatgtgccttcgggaaccgtgagacaggtgctgcatggctgtcgtcagctcgtgttgtgaaatgttgggttaagtcccgcaacgagcgcaacccttatcctttgttgccagcggtccggccgggaactcaaaggagactgccagtgataaactggaggaaggtggggatgacgtcaagtcatcatggcccttacgaccagggctaAaAaAgtgctacaatggcgcatacaaagagaagcgacctcgcgagagcaagcggacctcataaagtgcgtcgtagtccggattggagtctgcaactcgactccatgaagtcggaatcgctagtaatcgtggatcagaatgccacggtgaatacgttcccgggccttgtaca

m89 (5' to 3'):
tgacgggggcccgcacaagcggtggagcatgtggtttaattcgTTcgcgaagaaccttacctggtcttgacatccacggaagttttcagagatgagaatgtgccttcgggaaccgtgagacaggtgctgcatggctgtcgtcagctcgtgttgtgaaatgttgggttaagtcccgcaacgagcgcaacccttatcctttgttgccagcggtccggccgggaactcaaaggagactgccagtgataaactggaggaaggtggggatgacgtcaagtcatcatggcccttacgaccagggctacacacgtgctacaatggcgcatacaaagagaagcgacctcgcgagagcaagcggacctcataaagtgcgtcgtagtccggattggagtctgcaactcgactccatgaagtcggaatcgctagtaatcgtggatcagaatgccacggtgaatacgttcccgggccttgtaca

m90 (5' to 3'):
tgacgggggcccgcacaagcggtggagcatgtCCtttaattcgatgcaacgcgaagaaccttacctggtcttgacatccacggaagttttcagagatgagaatgtgccttcgggaaccgtgagacaggtgctgcatggctgtcgtcagctcgtgttgtgaaatgttgggttaagtcccgcaacgagcgcaacccttatcctttgttgccagcggtccggccgggaactcaaaggagactgccagtgataaactggaggaaggtggggatgacgtcaagtcatcatggcccttacgaccagggctaGaGacgtgctacaatggcgcatacaaagagaagcgacctcgcgagagcaagcggacctcataaagtgcgtcgtagtccggattggagtctgcaactcgactccatgaagtcggaatcgctagtaatcgtggatcagaatgccacggtgaatacgttcccgggccttgtaca

m91 (5' to 3'):
tgacgggggcccgcacTagcggtggagcatgtggtttaattcgatgcaacgcgaagaaccttacctggtcttgacatccacggaagttttcagagatgagaatgtgccttcgggaaccgtgagacaggtgctgcatggctgtcgtcagctcgtgttgtgaaatgttgggttaagtcccgcaacgagcgcaacccttatcctttgttgccagcggtccggccgggaactcaaaggagactgccagtgataaactggaggaaggtggggatgacgtcaagtcatcatggcccttacgaccagggctacacacgtgctacaatggcgcatacaaagagaagcgacctcgcgagagcaagcggacctcataaagtgcgtcgtagtccggattggagtctgcaactcgactccatgaagtcggaatcgctagtaatcgtggatcagaatgccacggtgaatacgttcccgggccttgtaca

m92 (5' to 3'):

tgacgggggcccCcTGaagcggtggagcatgtggtttaattcgatgcaacgcgaagaaccttacctggtcttgacatccacggaagttttcagagatgagaatgtgccttcgggaaccgtgagacaggtgctgcatggctgtcgtcagctcgtgttgtgaaatgttgggttaagtcccgcaacgagcgcaacccttatcctttgttgccagcggtccggccgggaactcaaaggagactgccagtgataaactggaggaaggtggggatgacgtcaagtcatcatggcccttacgaccagggctacacacgtgctacaatggcgcatacaaagagaagcgacctcgcgagagcaagcggacctcataaagtgcgtcgtagtccggattggagtctgcaactcgactccatgaagtcggaatcgctagtaatcgtggatcagaatgccacggtgaatacCAtccGgggccttgtaca

m93 (5' to 3'):
tgacgggggcccgcacaagcggtggagcatgtggtttaattcgatgcaacgcgaagaaccttacctggtcttgacatccacggaagttttcagagatgagaatgtgccttcgggaaccgtgagacaggtgctgcatggctgtcgtcagctcgtgttgtgaaatgttgggttaagtcccgcaacgagcgcaacccttatcctttgttgccagcggtccggccgggaactcaaaggagactgccagtgataaactggaggaaggtggggatgacgtcaagtcatcatggcccttacgaccagggctacacacgtgctacaatggcgcatacaaagagaagcgacctcgcgagagcaagcggacctcataaagtgcgtcgtagtccggattggagtctgcaactcgactccatgaagtcggaatcgctagtaatcgtggatcagaatgccacggtgaatacTTGTGcgggccttgtaca

m94 (5' to 3'):
tgacgggggcccgcacaagcggtggagcatgtggtttaattcgatgcaacgcgaagaaccttacctggtcttgacatccacggaagttttcagagatgagaatgtgccttcgggaaccgtgagacaggtgctgcatggctgtcgtcagctcgtgttgtgaaatgttgggttaagtcccgcaacgagcgcaacccttatcctttgttgccagcggtccggccgggaactcaaaggagactgccagtgataaactggaggaaggtggggatgacgtcaagtcatcatggcccttacgaccagggctacacacgtgctacaatggcgcatacaaagagaagcgacctcgcgagagcaagcggacctcataaagtgcgtcgtagtccggattggagtctgcaactcgactccatgaagtcggaatcgctagtaatcgtggatcagaatgccacgAAAAAAAAAAAAcgggccttgtaca

**Table S2: List of modified DNA oligos.**

| Oligonucleotide ID | DNA/RNA Sequence (5' to 3') |
|---|---|
| p0030_ab_fw | TCACGAAAGCTGAGTAGTCACGAGTCTTCT/idSp/CTGGCAGTTTTAGGCTGATTTGG |
| p0075_ab_bw | CCTTAATCATACTCACCAAATTACCATCCC/idSp/ATAAACGCAGAAAGGCCCAC |
| p0088_2xCy3.5 | Cy3p5-GGGATGGTAATTTGG[dT-Cy3p5]GAGTATGATTAAGG |
| p0109-biotin | 5BiotinTEG/TTATCCGCTCACAATTCCACA |
| p44 | GGTGTGTTGGTTGGGTGGTGGTAGT/idSp/GCAGGTCGACTCTAGAGGAT |
| p66-Cy3-FQ | /5Cy3/GGTGTATGTAATTGGAGTGGTT/3IABkFQ/ |
| P153-Cy3B-FQ | /5Cy3B/GGTGTATGTAATTGGAGTGGTT/3IABkFQ/ |
| P112A | 5BiotinTEG/AAAAAAcgcacaagAAA/3Cy3sp/ |
| prKG047 | /5Cy3/AAAugugguuuAAAAAA/3biotinTEG/ |
| prKG078 | 5BiotinTEG/AAAAAAccuggucuAAA/3Cy3sp/ |
| prKG081 | 5BiotinTEG/AAAAAAcctggtctAAA/3Cy3sp/ |
| p110Bcy5 | CTTGTGC/3Cy5sp/ |
| prKG058cy5.5 | CTTGTGC/3Cy55sp/ |
| prKG080cy7 | CTTGTGC/3Cy7p/ |
| prKG032cy5 | /5Cy5/AACCACA |
| prKG057cy7 | /5Cy7/AACCACA |

| | |
|---|---|
| prKG053cy5 | AGACCAG/3Cy5sp/ |
| prKG072 | TGACGACAGCCATGCAGC |
| prKG073 | GTAGCCCTGGTCGTAAGG |
| prKG074 | GTTCTTCGCGTTGCATCG |
| prKG075 | TGTATGCGCCATTGTAGC |
| prKG076 | CGATTACTAGCGATTCCG |
| prKG077 | TGTACAAGGCCCGGGAAC |

/idSp/ denotes abasic site.
[dT-Cy3p5] is an internal modification labeled on the nucleobase of deoxythymidine.
5BiotinTEG denotes Biotin-TEG attached to 5'-end
3BiotinTEG denotes Biotin-TEG attached to 3'-end
/3IABkFQ/ denotes FQ quencher attached to 3'-end
/5Cy3/ denotes Cy3 label attached to 5'-end
/5Cy3B/ denotes Cy3B label attached to 5'-end
/3Cy3sp/ denotes Cy3 label attached to 3'-end
/5Cy5/ denotes Cy5 label attached to 5'-end
/3Cy5sp/ denotes Cy5 label attached to 3'-end
/3Cy55Sp/ denotes Cy5.5 label attached to 3'-end
/3Cy7p/ denotes Cy7 label attached to 3'-end
/5Cy7/ denotes Cy7 label attached to 5'-end


1. O. Duss, G. A. Stepanyuk, J. D. Puglisi, J. R. Williamson, Transient Protein-RNA Interactions Guide Nascent Ribosomal RNA Folding. *Cell* **179**, 1357-1369 e1316 (2019).
2. M. Kitagawa *et al.*, Complete set of ORF clones of Escherichia coli ASKA library (a complete set of E. coli K-12 ORF archive): unique resources for biological research. *DNA Res* **12**, 291-299 (2005).
3. O. Duss *et al.*, Real-time assembly of ribonucleoprotein complexes on nascent RNA transcripts. *Nat Commun* **9**, 5087 (2018).
4. N. S. Qureshi, O. Duss, Tracking transcription-translation coupling in real time. *Nature*, (2024).
5. S. D. Chandradoss *et al.*, Surface passivation for single-molecule protein studies. *J Vis Exp*, (2014).
6. C. E. Aitken, R. A. Marshall, J. D. Puglisi, An oxygen scavenging system for improvement of dye stability in single-molecule fluorescence experiments. *Biophys J* **94**, 1826-1835 (2008).
7. I. Rasnik, S. A. McKinney, T. Ha, Nonblinking and long-lasting single-molecule fluorescence imaging. *Nat Methods* **3**, 891-893 (2006).
8. M. F. Juette *et al.*, Single-molecule imaging of non-equilibrium molecular ensembles on the millisecond timescale. *Nat Methods* **13**, 341-344 (2016).
9. J. Chen *et al.*, High-throughput platform for real-time monitoring of biological processes by multicolor single-molecule fluorescence. *Proc Natl Acad Sci U S A* **111**, 664-669 (2014).
10. J. Chen, A. Petrov, A. Tsai, S. E. O'Leary, J. D. Puglisi, Coordinated conformational and compositional dynamics drive ribosome translocation. *Nat Struct Mol Biol* **20**, 718-727 (2013).
